# Modulating immune cell fate and inflammation through CRISPR-mediated DNA methylation editing

**DOI:** 10.1101/2024.07.10.599183

**Authors:** Gemma Valcárcel, Anna V. López-Rubio, Aleksey Lazarenkov, Clara Berenguer, Josep Calafell, Javier Rodríguez-Ubreva, Esteban Ballestar, José Luis Sardina

## Abstract

DNA methylation is traditionally associated with gene silencing, but its causal relationship and role in shaping cell fate decisions still need to be fully elucidated. Here, we conducted a genome-wide analysis to investigate the relationship between DNA methylation and gene <u>expression</u> at gene regulatory regions in human immune cells. By utilizing CRISPR-dCas9 DNA methylation editing tools, we successfully established a cause-and-effect relationship between the methylation levels of the promoter of the Interleukin1-receptor antagonist (*IL1RN*) gene and its expression. Notably, we observed that modifying the DNA methylation status of the *IL1RN* promoter is sufficient to alter the acquisition of the human myeloid cell fate and change the cellular response to inflammatory stimuli, resulting in abnormal cytokine release and distinctive capacity to support cancer growth.

## INTRODUCTION

DNA methylation (DNAm) at CpG residues is the main epigenetic modification of mammalian DNA and is commonly associated with transcriptional repression^1^. This link arises from extensive research in the fields of differentiation and development, investigating the effects of disrupting the function of key DNA modifiers such as DNMT and TET enzymes on transcription^2–4^. Alternatively, in cancer research, drugs such as azacitidine and decitabine, which are DNA hypomethylating agents, have been widely used to investigate these connections^5,6^. Both approaches offer valuable insights, but they have the potential to alter the overall methylome of the cells, posing a challenge in distinguishing between direct and indirect effects on gene expression. DNMTs and TETs, in addition to their catalytic activity, can interact with transcription- and chromatin-related factors, thus impacting transcription independently of DNAm^7–12^. On top of that, these a priori antagonistic enzymes have been described as requiring each other to develop their genome-wide catalytic activity properly^13,14^, adding another layer of complexity to the interpretation of the loss-of-function experiments. Finally, hypomethylating small molecules have diverse biological functions unrelated to DNAm^15^. Therefore, the extent to which DNAm directly instructs gene expression remains unclear.

A better understanding of the instructive role of DNAm in transcriptional outcomes was achieved with the development of tools that allow precise addition or removal of DNAm marks^6,16^. For instance, by employing a modular CRISPR-Cas9-based epigenome editing tool coupled with a single-cell and quantitative readout, Policarpi and co-workers showed DNAm, leading to penetrant transcriptional silencing^17^. However, this DNAm-induced silencing was not as strong as the repression associated with other epigenetic marks, such as H2AK119ub or H3K9me2/3^17^. Genome-wide studies have revealed context-specific transcriptional responses associated with DNAm gain at promoter regions, including increased gene expression^18^. The diverse responses may be influenced by the varying sensitivity of different transcription factor families to the presence of DNAm at their specific recognition sites in the genome^19^. Additionally, recent evidence has shown that DNAm can cause transcriptional activation by counteracting the polycomb-mediated deposition of H3K27me3^20^.

The interplay between transcription factors and epigenetic regulators is essential during hematopoiesis, with DNAm playing a significant role in influencing immune cell identity and function^21^. However, the causal connections between DNAm and immune cell fates have not been thoroughly explored. In 2016, Amabile and co-workers demonstrated the stable editing of DNAm at multiple genomic locations and its connection to transcriptional silencing for the first time in immune cells^22^. More recently, a CRISPR-Cas9 DNAm editor was used to silence the *CDKN2B* gene in human hematopoietic stem cells, leading to a bias in the differentiation output from the edited cells^23^. Despite the importance of these reports, a fuller knowledge of the molecular mechanisms underlying altered hematopoiesis in response to DNAm editing is necessary due to its significant implications for differentiation and cancer. Furthermore, studies addressing the causal relationship between DNAm and relevant innate immune responses, such as inflammation, are lacking.

In our study, we investigated how changes in DNAm at gene regulatory regions impact gene expression during the conversion of human leukemic B cells into non-tumorigenic macrophages^24^. We employed CRISPR/dCas9 methylation editing technology to precisely alter the DNAm status of the *IL1RN* promoter and evaluate its impact on relevant immune cellular phenotypes. Our findings revealed that *IL1RN* plays a significant role in the acquisition of the myeloid cell fate in humans. Additionally, we noticed that the methylation-edited cells exhibited modified responses to inflammatory stimuli via the IL1 signaling pathway.

## RESULTS

### WGBS-seq uncovers the genome-wide reshaping of DNAm during human B to-macrophage transdifferentiation

To gain insight into the dynamics of DNAm during myeloid cell fate acquisition, we are utilizing the highly efficient and homogenous C/EBPα-driven conversion of human B leukemic cells containing a β-estradiol inducible form of C/EBPα (BlaER cells), into non-tumorigenic induced macrophages (iMacs)^24^. The iMacs generated using this protocol closely resemble their natural counterparts as they are fully phagocytic and have inflammasome competency^24–26^.

Cultured B cells were treated with β-estradiol (E2) and collected at different time points of the cell fate conversion process (0h -B cells-; 24h; 96h and 168h -iMacs-) **(Fig. 1a)**. We then generated genome-wide, nucleotide-resolution maps for DNAm by whole-genome bisulfite sequencing (WGBS-seq). DNAm levels were determined by computing the values of approximately 24 million CpG residues that were covered ≥5X in all the samples. Our B cells exhibited a widespread pattern of DNA hypermethylation, similar to what was observed in primary human B lymphocytes^27^. After induction of C/EBPα, a genome-wide reshaping of DNAm was observed from 96 hours onwards **(Fig. 1b and Extended Data Fig. 1a)**. Approximately 14% of the DNAm signal, which accounts for 384,127 1kb bins, was redistributed during this process **(Extended Data Fig. 1b)**. Interestingly, the redistribution of the DNAm signal did not occur randomly in the genome. Instead, it occurred preferentially on specific chromosomes, with ChrX showing the largest changes **(Extended Data Fig. 1b)**.

**Figure 1.**
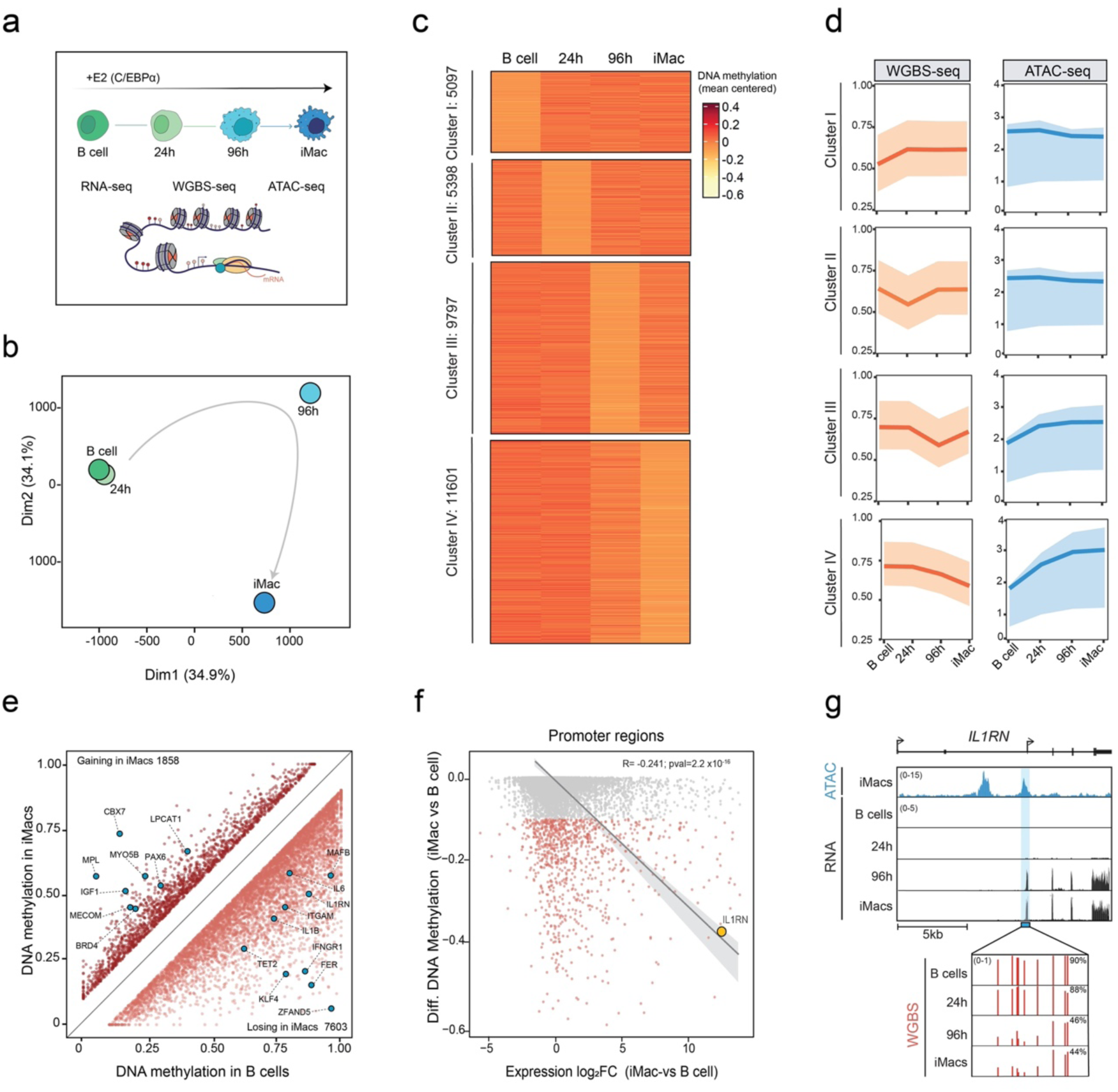
Epigenome profiling and integrative analyses uncover the *IL1RN* promoter as a top event showing a DNA methylation-expression correlation. **(a)** Schematic overview of samples and methodology used in the epigenome profiling. iMac: induced macrophages. **(b)** Multidimensional Scaling (MDS) analysis showing DNAm dynamics at 1kb bins genome-wide during transdifferentiation. A grey arrow indicates the hypothetical trajectory. **(c)** Clustering of genome-wide DNAm dynamics (by WGBS-seq) at 1kb bins containing chromatin accessible regions (ATAC+) associated with promoters (based on TSS proximity) or enhancers (based on H3K4me1 peaks) during transdifferentiation^26^. Only 1kb bins showing at least 10% DNAm changes are depicted. **(d)** Quantification of DNAm (left panels) and ATAC (right panels) signals at the clusters in (c). Plots represent the mean (line) and interquartile range (shaded region). ATAC-seq data from GSE131620^26,28^.**(e)** Scatter plot comparing DNAm levels at iMac’s chromatin accessible regions (ATAC+ peaks) in iMacs versus in B cells. Dark red dots indicate ATAC+ regions gaining at least 10% (n=1858) of DNAm in iMacs; light red dots indicate ATAC+ regions losing at least 10% (n=7603) of DNAm in iMacs. Blue dots highlight ATAC+ regions of interest. **(f)** Plot showing the correlation between the step changes (iMac-Bcells) in DNAm and gene expression at iMac’s chromatin accessible regions (ATAC+ peaks) overlapping annotated promoters. Grey dots: ATAC+ promoter regions not losing at least 10% in DNAm. Light red dots: ATAC+ promoter regions losing at least 10% of DNAm in iMacs. The yellow dot highlights the correlation between DNAm and expression at the *IL1RN* promoter region. RNA-seq data from GSE140528^26,28^. **(g)** Genome browser snapshot showing signal for chromatin accessibility (by ATAC-seq) in iMacs, and gene expression (by RNA-seq) during transdifferentiation at the *IL1RN* locus. The blue-shaded region represents the promoter of the *IL1RN* short isoform (ENST00000409930.4), which showed DNA demethylation (by WGBS-seq) during transdifferentiation.

These observations suggest that the chromatin environment might impact the reshaping of the DNAm signal during B to macrophage cell fate conversion.

### Myeloid-related Gene Regulatory Elements (GREs) are demethylated during macrophage cell fate acquisition

To further understand the genomic context where the redistribution of DNAm occurs, we investigated the interplay between DNAm and chromatin accessibility using ATAC-seq data^26,28^. To that end, we intersected the DNAm dynamic bins (**Extended Data Fig.1b**) with ATAC-seq data and selected those containing at least one peak of ATAC associated with a TSS (promoters) or with an H3K4me1 peak (enhancers)^26^. As a result, 31,893 DNAm dynamic bins containing Gene Regulatory Elements (GREs) were selected, accounting for approximately only 8% of all dynamic bins observed. This underscores that the majority of DNAm reshaping occurs distantly from GREs, potentially playing a limited role in the biology of this specific cell fate process. Therefore, we decided to focus all our further DNAm analyses exclusively on regions containing GREs. To achieve this, the approximately 32,000 dynamic DNAm bins containing GREs were grouped into 4 major clusters (Clusters I to IV) based on the timing of their DNAm changes. The clusters exhibited predominantly sequential and mostly transient changes in DNAm (clusters I to III), except for cluster IV **(Fig. 1c)**. Cluster IV, containing approximately one-third of the total number of regions, showed continuous loss of DNAm from 96 hours onward **(Fig. 1c)**. Of note, only regions included in clusters showing demethylation from 96 hours onwards, regardless of transient or continuous behavior, experienced an increase in chromatin accessibility (by ATAC-seq) **(Fig. 1d and Extended Data Fig. 1c-d)**. Conversely, no chromatin closure was observed in association with gain of DNAm (cluster I regions) **(Fig.1d and Extended Data Fig. 1c-d).** Thus suggesting DNAm gain might not regulate chromatin activity in this cellular model of myeloid cell fate acquisition. To further investigate this hypothesis, we specifically analyze changes in DNAm at chromatin-accessible regions (ATAC-positive) between B cells and iMacs. In iMacs, we observed 7,603 ATAC-positive regions that lost DNAm and 1,858 regions that gained it, showing preferential association with Clusters III and IV, and Cluster I, respectively **(Fig. 1e and Extended Data Fig. 1e)**. The regions of DNAm loss and gain exhibited different genomic distributions **(Extended Data Fig. 1f)**. The gain regions mainly overlapped with introns, and promoter regions at similar percentages (35.9% and 29.1%, respectively). Conversely, in the loss regions, there was a preferential enrichment in intronic regions, possibly intragenic enhancers, at the expense of promoters (41.8% and 21%, respectively) **(Extended Data Fig. 1f)**. To uncover biological processes potentially regulated by the DNAm changes described, we performed Gene Ontology (GO) analysis at the ATAC-positive DNAm loss and gain regions **(Extended Data Fig. 1g)**. This showed enrichment of terms related to myeloid cells for genes associated with loss regions, including “Myeloid leukocyte migration,” “Phagocytosis,” and “Positive regulation of cytokine production,” among others. On the other hand, the gain regions were unspecifically associated with terms related to organ development, such as “Axon guidance,” “Kidney development,” and “Heart morphogenesis,” among others **(Extended Data Fig. 1g)**. In line with these findings, TF motif analysis showed significant enrichment of myeloid transcription factors, such as AP1 complex subunits (including Fos, JunB, BATF, Fra1-2, among others) and CEBP factors, in the ATAC-positive DNAm loss regions **(Extended Data Fig. 1h)**. In contrast, the DNAm gain regions displayed a modest enrichment in the motifs of genome architecture factors, including CTCF and its relative BORIS (also known as CTCFL), along with development-related transcription factors such as Hox, Slug, or Snail, among others **(Extended Data Fig. 1h)**.

Altogether, this suggests that specific GREs related to myeloid cells are undergoing DNA demethylation and potential chromatin activation during the cell fate conversion process. At the same time, GREs unrelated to the cell fate conversion process are gaining DNAm without apparent changes in chromatin activity.

### Integrative analyses reveal the *IL1RN* promoter as a top event showing a DNA methylation-expression correlation during transdifferentiation

To gain mechanistic insight into the transcriptional consequences of the DNAm changes observed at GREs, we investigated the interplay between DNAm and gene expression using RNA-seq data^26,28^. We observed no correlation between the gain of DNAm at ATAC-positive regions and changes in gene expression of their associated genes (R= -0.018; pval= 0.64) **(Extended Data Fig. 1i)**. Meanwhile, a significant negative correlation was detected between DNAm loss and gene expression, both at ATAC-positive regions non-associated with promoters (R= -0.132; pval=7.2x10^-10^) **(Extended Data Fig.1j)** and an even stronger correlation when they were overlapping promoters (R= -0.241; pval=2.2x10^-16^) **(Fig. 1f)**. The promoter of the Interleukin-1 receptor antagonist (*IL1RN*) gene, which shows a DNAm loss over 40% and an expression increase of more than 1000-fold between iMacs and B cells, emerged as a ltop correlated event in the analysis **(Fig. 1f)**. The *IL1RN* gene, which plays an essential role in inflammatory responses in mature myeloid cells^29^, is not expressed and its promoter is fully methylated at the B cell stage **(Fig. 1g)**. However, the methylation of the *IL1RN* promoter decreases between 24 and 96 hours, which coincides with the reactivation of the gene **(Fig. 1g)**. These changes occur specifically in the promoter of the *IL1RN* short isoform, IL1RN-205 (ENST00000409930.4), which corresponds to the secretory protein variant preferentially expressed in myeloid cells^29^.

These analyses suggest that only gene regulatory regions losing DNAm, such as the promoter of the *IL1RN* gene, but not regions gaining it, are correlated with changes in gene expression during transdifferentiation.

### dCas9-TET1-mediated demethylation of the *IL1RN* promoter is sufficient to lead to its gene reactivation

While the phenotypes associated with the loss of function of major DNAm regulators have been extensively studied, the impact of individual methylation events on immune phenotypes remains largely unexplored, mainly due to technical limitations. This has been recently challenged with the advent of DNAm editing tools, especially those associated with CRISPR technology^30^.

Since the *IL1RN* promoter is hypermethylated and its associated gene is not expressed in B cells **(Fig. 1e-g)**, we speculated about the possibility of inducing DNA demethylation specifically at this region to force *IL1RN* expression at the B cell stage. To address this challenge, we stably integrated a previously developed DNA demethylation editing tool^31,32^, into our human B leukemic cells **(Fig. 2a)**. The editing tool is based on a catalytically dead Cas9 protein tethered with the catalytic domain of the DNA dioxygenase TET1 (dCas9-TET1). B cell clones expressing high levels of the dCas9-TET1 transgene were selected and expanded. Afterward, we transduced dCas9-TET1 clones with lentiviruses simultaneously encoding 4 different sgRNAs that targeted either no sequence within the genome (CTRL cells) or the *IL1RN* promoter (sgIL1RN cells) **(Fig. 2a)**. We confirmed that the dCas9-TET1 protein was specifically bound to the *IL1RN* promoter in sgIL1RN B cells **(Fig. 2b)**. The dCas9-TET1 recruitment led to the demethylation of important CpG residues located upstream of the gene’s TSS **(Fig. 2c)**. Among them, the largest demethylation (approximately 50% decrease) was observed at the CpG residue located just upstream of the TSS (-9bp) **(Fig. 2c)**. Finally, because of the DNA demethylation mediated by dCas9-TET1 we detected *IL1RN* gene activation (>15-fold increase in mRNA levels) and protein accumulation (>60-fold increase) in sgRNA B cells **(Fig. 2d)**.

**Figure 2.**
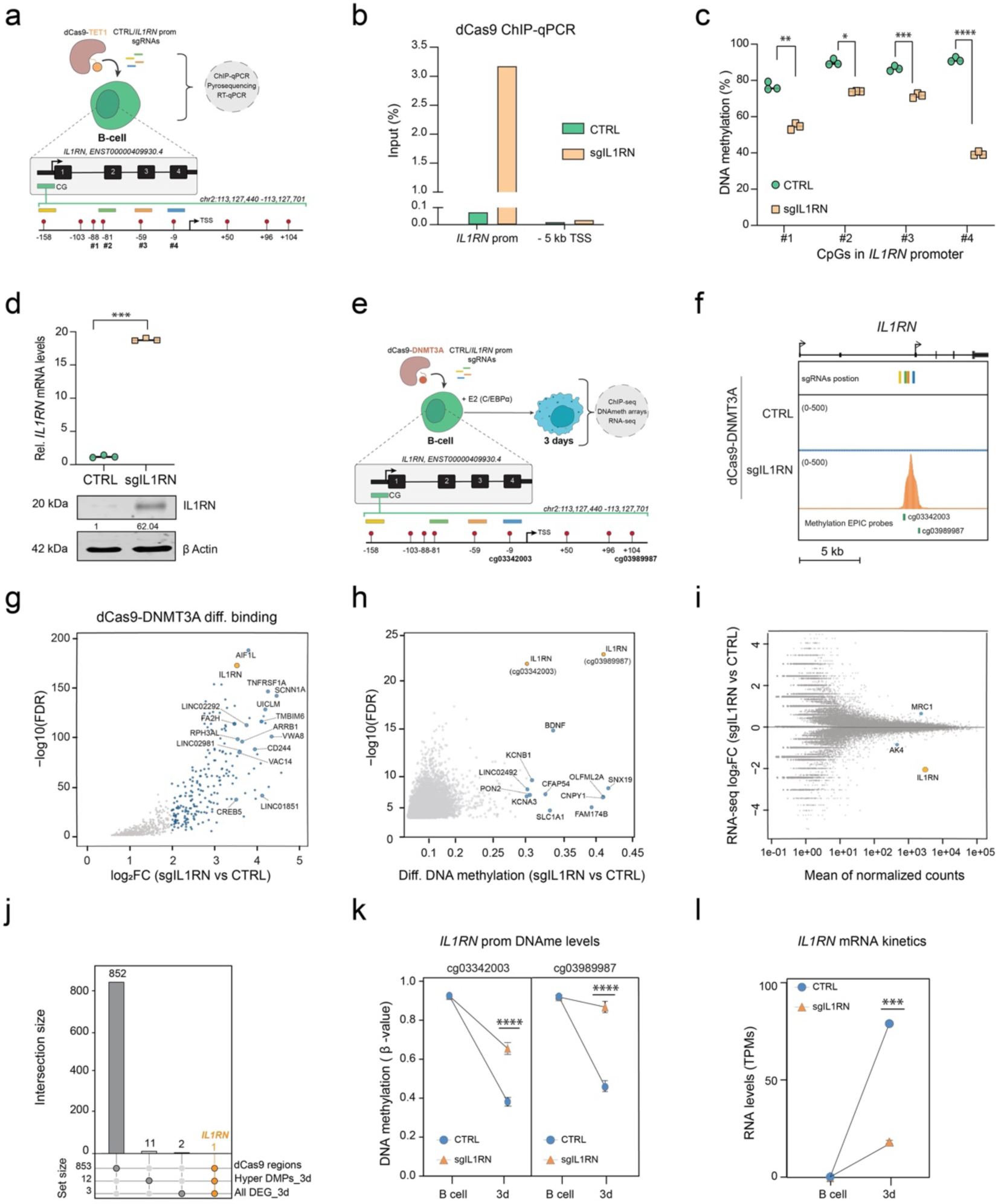
CRISPR-mediated epigenome editing at the *IL1RN* promoter efficiently modulates DNAm levels and gene expression. **(a)** Schematic of the dCas9-TET1 *IL1RN* promoter epigenome editing experiment. Top, lentiviral transduction was used to introduce the dCas9-TET1 epigenome editing tool into B cells, along with sgRNAs that targeted either no sequence within the genome (CTRL) or the *IL1RN* promoter (sgIL1RN). Bottom, diagram representation of the four sgRNAs selected targeting regions upstream the transcription start site (TSS) of the *IL1RN* gene. The sgRNAs recognizing their respective target site are shown in yellow, green, orange, and blue, with the CpG sites as red lollipops. #1 to #4 indicate the CpG residues analyzed in the experiment **(b)** ChIP-qPCR showing the enrichment of dCas9-TET1 at the *IL1RN* prom region in dCas9-TET1 sgIL1RN and CTRL B cells. A region 5kb downstream of the *IL1RN* prom is shown as a negative control region. **(c)** DNAm levels (by pyrosequencing) at 4 CpG residues located upstream of the *IL1RN* TSS (as shown in a) in dCas9-TET1 sgIL1RN and CTRL B cells. One-way ANOVA with Dunnett’s post-hoc correction, n=3, mean ± s.e.m., (*p< 0.05; **p< 0.01, ***p< 0.001,****p< 0.0001). **(d)** Top: *IL1RN* Expression (RT-qPCR) in dCas9-TET1 sgIL1RN and CTRL B cells. Values were normalized against *HPRT* expression. Unpaired two-tailed Student’s t-test, n=3, mean s.e.m., (***p< 0.001) Bottom: Representative western blot image of IL1RN protein in dCas9-TET1 sgIL1RN and CTRL B cells. IL1RN levels were normalized to b-Actin levels and expressed as a fold change over CTRL B cells. **(e)** Schematic of the dCas9-DNMT3A *IL1RN* promoter epigenome editing experiment. Top, lentiviral transduction was used to introduce the dCas9-DNMT3A epigenome editing tool into B cells, along with sgRNAs that targeted either no sequence within the genome (CTRL) or the *IL1RN* promoter (sgIL1RN). Bottom, as explained in (a). 2 Infinium MethylationEPIC v2.0 probes are shown in –9 (cg03342003) and +104 (cg03989987) positions. **(f)** Genome browser snapshot showing the dCas9-DNMT3A binding at the *IL1RN* promoter region. Cas9 ChIP-seq profiles are shown for dCas9-DNMT3A CTRL and sgIL1RN B cells. The location of the 4 sgRNAs targeting *the IL1RN* promoter and the 2 array probes is depicted. **(g)** Scatter plot showing the genome-wide differential dCas9-DNMT3A enrichment in sgIL1RN B cells over CTRL B cells. Blue dots indicate regions significantly more enriched in dCas9-DNMT3A in sgIL1RN B cells than CTRL B cells (FC>2, p<0.05, n=853). Large blue dots and the yellow dot (*IL1RN* prom) indicate top dCas9-DNMT3A bound promoter regions (FC>3.5, p<0.05) that will be further analyzed. **(h)** Scatter plot showing differentially hypermethylated CpGs in sgIL1RN day3 cells compared to CTRL day3 cells. Grey dots indicate significantly hypermethylated CpG positions (FDR < 0.05). Blue and yellow dots indicate significantly hypermethylated CpG positions gaining ≥ 30% of DNAmth (FDR < 0.05, Δβ ≥ 0.3, n=13). The yellow dots highlight two differentially hypermethylated CpG positions in the *IL1RN* prom (probes cg03342003 and cg03989987 from the MethylationEPIC BeadChip 850k v2.0 microarrays) **(i)** MA plot showing differentially expressed genes (DEGs) in sgIL1RN day3 cells compared to CTRL day3 cells. Blue dots and yellow dot (*IL1RN*) indicate the significant DEGs (FC >1, p<0.05, n=3). **(j)** Upset plot depicting the intersection of the significantly dCas9-DNMT3A bound regions (in f), the significantly hypermethylated Δβ ≥ 0.3 CpG positions (in g), and the significant associated DEGs (in h). *IL1RN* is the only candidate intersecting the 3 datasets. **(k)** DNA methylation dynamics (array data) of two differentially hypermethylated CpG positions (probes cg03342003 and cg03989987) at the *IL1RN* prom in sgIL1RN and CTRL cells. Unpaired two-tailed Student’s t-test, n=4, mean s.e.m., (****p< 0.0001). **(l)** *IL1RN* expression dynamics (by RNA-seq) in sgIL1RN and CTRL cells. Unpaired two-tailed Student’s t-test, n=2, mean s.e.m., (***p< 0.001).

Here, we utilized the dCas9-TET1 editing tool to successfully activate the expression of a myeloid gene in an inappropriate immune cell type (B-lymphoid cells). This allowed us to unveil a causal relationship between the DNAm status of the *IL1RN* promoter and the expression of its gene.

### dCas9-DNMT3A efficiently and specifically hypermethylates the *IL1RN* promoter, leading to its downregulation

As previously shown, the *IL1RN* promoter is demethylated between 24 and 96 hours of the cell fate conversion process, coinciding with the reactivation of its gene **(Fig. 1g)**. Therefore, based on the observed causality between DNAm and *IL1RN* expression, we hypothesized that blocking the DNAm loss associated with the transdifferentiation process would alter *IL1RN* gene expression patterns **(Fig. 2c-d)**. To tackle this task, we introduced a previously developed DNAm editing tool^31,32^, into our human B leukemic cells using lentiviral infection **(Fig. 2e)**. The editing tool is based on a catalytically dead Cas9 protein tethered with the catalytic domain of the DNA de novo methyltransferase DNMT3A (dCas9-DNMT3A). dCas9-DNMT3A-expressing stable clones were selected, expanded, and transduced with lentiviruses encoding 4 different sgRNAs targeting either no sequence within the genome (CTRL) or the *IL1RN* promoter (sgIL1RN) **(Fig. 2e)**. We confirmed that the dCas9-DNMT3A protein was strongly and specifically bound to the *IL1RN* promoter in sgIL1RN B cells **(Fig. 2f and Extended Data Fig. 2a)**. However, as expected from previous research using these DNAm editor tools^32–34^, we also observed the off-target binding of the dCas9-DNMT3A protein to 852 genomic regions **(Fig. 2g)**. Of note, some of the strongest off-target sites (FC>3.5) were identified as overlapping promoters. Therefore, the recruitment of the dCas9-DNMT3A protein may result in the hypermethylation of these promoters and the deregulation of their genes. However, in B cells, none of the off-target regions or the *IL1RN* promoter itself (that is already hypermethylated) showed differential methylation and their associated genes were not differentially expressed **(Extended Data Fig. 2b-f)**. We then examined the situation 3 days after C/EBPα activation. By this time, the *IL1RN* promoter should have started to lose DNAm, allowing us to detect differences in their methylation levels between CTRL and sgIL1RN cells. We found out that only 13 CpG residues were hypermethylated, with two of them displaying the most significant changes in the *IL1RN* promoter **(Fig. 2h)**. Additionally, we observed only 3 Differentially Expressed Genes (DEGs), with *IL1RN* being downregulated more than 4-fold in sgIL1RN 3d cells **(Fig. 2i)**.

Overall, the only region differentially occupied by dCas9-DNMT3A, differentially methylated and differentially expressed when comparing sgIL1RN cells and CTRL cells is the *IL1RN* gene **(Fig. 2j-l and Extended Data Fig. 2f-g)**. Thus, the specificity and accuracy of the dCas9-DNMT3A-mediated DNAm editing were validated, and the previously described causality between DNAm and expression at this gene was reinforced.

### dCas9-DNMT3A-mediated hypermethylation of the *IL1RN* promoter leads to impaired human myeloid cell fate acquisition and enhanced phagocytic capacity

The IL1 pathway has been described as playing a role in maintaining balanced hematopoiesis between the lymphoid and myeloid lineages in mice. Both chronic exposure to IL1β^35^ and genetic deficiency of *Il1rn*^36^ have been described as leading to hyperactivation of the IL1 pathway, which causes a bias towards the myeloid lineage. The exact role of IL1RN and the IL1 pathway in the acquisition of human myeloid cell fate, as well as the molecular mechanisms underlying the myeloid bias following hyperactivation of the IL1 pathway, are not yet fully understood.

The kinetics of *IL1RN* expression during our human B-to-macrophage transdifferentiation system suggest a potential involvement of IL1RN and the IL1 pathway in the physiological acquisition of the human myeloid cell fate. To explore this possibility, we analyzed the transcriptomic profiles of CTRL and sgILRN cells throughout the transdifferentiation process. As previously observed, minor changes in gene expression were detected at short time points **(Extended Data Fig. 3a)**. This is expected since *IL1RN* expression is detected from day 3 onwards **(Fig. 2i,l and Extended Data Fig. 2e)**. However, at the iMac stage, a distinct transcriptional profile emerged between the CTRL and sgILRN cells **(Extended Data Fig. 3a-b)**. Approximately 2,000 DEGs were detected at this stage. Among these, 912 genes were up-regulated, and 1002 were down-regulated. One of the down-regulated genes was *IL1RN* itself, which showed a strong reduction of about 16-fold in its mRNA levels **(Extended Data Fig. 3b)** and was largely decreased at the protein level in dCas9-DNMT3A sgIL1RN iMacs **(Extended Data Fig. 3c)**. To further investigate the impact of *IL1RN* DNAm editing on the transcriptomic kinetics, all the genes differentially expressed in dCas9-DNMT3A CTRL and sgILRN cells throughout the transdifferentiation process were selected and clusterized into 4 major groups (Transdifferentiation Clusters -TC- 1 to 4) **(Fig. 3a)**. As expected, differences in gene expression within the clusters between CTRL and sgIL1RN cells were only detectable at the iMac stage **(Fig. 3a-b)**. In TC1, the *IL1RN* DNAm editing led to an incomplete silencing of genes related to cellular division, such as the well-known *BRCA1* or *CDK1* genes **(Fig. 3b-c)**. In TC2, IL1RN DNAm editing further enhanced the downregulation of genes involved in the transcriptional program of B cells, including the genes coding for the B cell transcription factor *LEF1* and the B cell marker *CD79A* **(Fig. 3b-c)**. In TC3, the DNAm editing prevented the decrease in gene expression for a group of transiently upregulated genes related to metabolic processes, including Hexokinase 2 (*HK2*) and Phosphoglycerate kinase 1 (*PGK1*) **(Fig. 3b-c)**. Finally, in TC4, the DNAm editing hindered the upregulation of genes related to myeloid cell fate and phagocytosis processes, including *ITGAM* and *CD14* **(Fig. 3b-c)**. Altogether, these transcriptional analyses suggest that modifying the methylation status of the *IL1RN* promoter could modify the cell fate acquired during the transdifferentiation process, leading to cells with altered cell proliferation, metabolism, and myeloid differentiation status. Notably, this notion was supported by the observation that *IL1RN* DNAm edited iMacs showed reduced levels of the typical myeloid markers CD11b (encoded by *ITGAM*) and CD14 **(Fig. 3d)** and an enhanced phagocytic capacity **(Fig. 3e)**.

**Figure 3.**
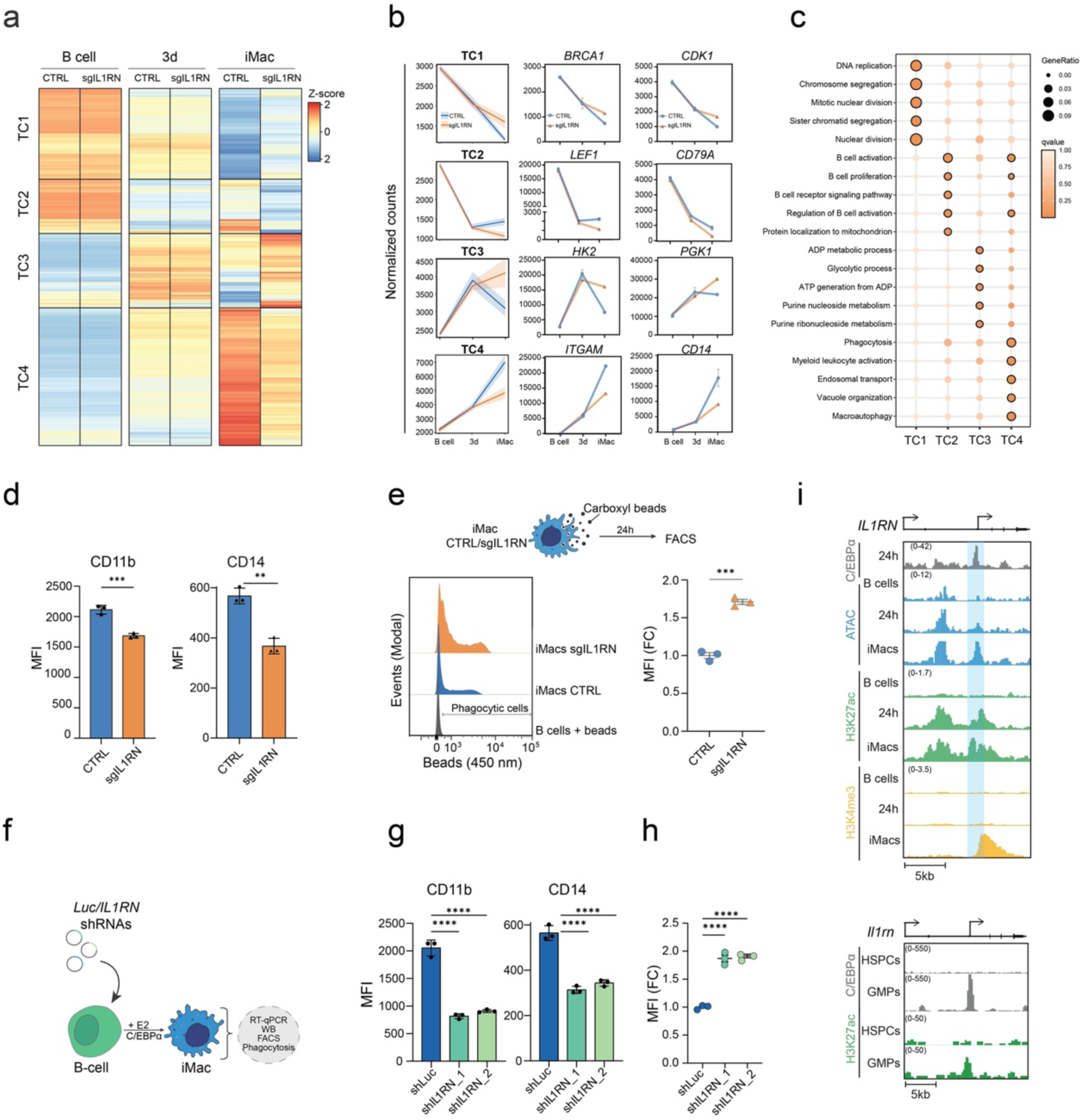
*IL1RN* DNAm editing alters the acquisition of the myeloid cell fate and the phagocytic capacity. **(a)** Clustering of the transcriptional dynamics (by RNA-seq) during transdifferentiation in sgIL1RN and CTRL cells. Average scaled normalized counts (Z-score) of differentially expressed genes (DEG) (padj < 0.05) are represented at each timepoint (n=2 biologically independent replicates). Black lines show the splitting by k-means clustering. Transdifferentiation Clusters (TC1-4). **(b)** Left panel: Quantification of RNA signal at the transdifferentiation clusters (in a) in sgIL1RN and CTRL cells.; Center and right panels: expression dynamics of representative genes from the clusters. **(c)** Balloon plot depicting the Gene Ontology (GO) enrichment analysis for the genes associated with clusters in a. The top 5 most significantly over-represented biological processes (BP) terms for each cluster are plotted. Ratio of genes of interest over all unique genes (GeneRatio) and qvalue are shown. Significant terms (qvalue < 0.05) are highlighted with a black stroke. **(d)** Flow cytometric analysis showing the Mean Fluorescence Intensity (MFI) for the myeloid markers (CD11b and CD14) in sgIL1RN and CTRL iMacs. Unpaired two-tailed Student’s t-test, n=3, mean s.e.m., (**p< 0.01; ***p< 0.001). **(e)** Flow cytometric analysis showing the phagocytic capacity of sgIL1RN and CTRL iMacs. Left: Representative experiment showing the uptake of blue fluorescent beads. Right: Plot showing the normalized Mean Fluorescence Intensity (MFI) for the blue beads in sgIL1RN and CTRL iMacs. Unpaired two-tailed Student’s t-test, n=3, mean s.e.m., (***p< 0.001) **(f)** Schematic of the *IL1RN* shRNAs experiments during transdifferentiation. B cells were transduced with lentiviruses encoding shRNAs targeting the luciferase gene (shLuc, control) or the *IL1RN* gene (shIL1RN). After puromycin selection, cells were treated for 7 days with E2 (β-estradiol) to induce transdifferentiation from B cells to iMacs. **(g)** Flow cytometric analysis showing the Mean Fluorescence Intensity (MFI) for the myeloid markers (CD11b and CD14) in shIL1RN and shLuc iMacs. Two-way ANOVA with Dunnett’s post-hoc test, n=3, mean ± s.e.m., (****p < 0.0001). **(h)** Plot showing the normalized Mean Fluorescence Intensity (MFI) for the blue beads in shIL1RN and shLuc iMacs. Two-way ANOVA with Dunnett’s post-hoc test, n=3, mean ± s.e.m., (****p < 0.0001). **(i) Top**: Genome browser snapshot at the human *IL1RN* locus showing signal for C/EBPα ChIP-seq at 24h and chromatin accessibility (by ATAC-seq), H3K27ac and H3K4me3 ChIP-seq signal in B cells, 24 hours and in iMacs. The blue-shaded region indicates the DNA demethylated region at the *IL1RN* promoter (as in Figure 1E-G). **Bottom:** Genome browser snapshot at the mouse *Il1rn* locus showing signal for C/EBPα and H3K27ac ChIP-seq in murine Hematopoietic Stem Progenitor cells (HSPCs) and Granulo-Monocyte Progenitors (GMPs). Data were taken from: H3K27ac and H3K4me3 ChIP-seq during human transdifferentiation (ArrayExpress: E-MTAB-9010); C/EBPα ChIP-seq in murine HSPCs and GMPs (GEO: GSE43007)^71^ H3K27ac ChIP-seq signal in murine HSPCs and GMPs (GEO: GSE59636)^72^.

### An orthogonal approach using shRNAs validates the involvement of IL1RN in human myeloid cell fate acquisition and phagocytic activity

To validate our relevant results obtained with the dCas9-DNMT3A tool, we employed an orthogonal method. We infected our human B leukemic cells with lentiviruses harboring shRNAs that target the *IL1RN* mRNA. We used shRNAs against the firefly luciferase (Luc) as a control **(Fig. 3f)**. Both shRNAs efficiently depleted IL1RN at the mRNA and protein levels in iMacs **(Extended Data Fig. 3d-e)**. The *IL1RN*-depleted (shIL1RN) iMacs displayed similar changes in gene expression as observed in *IL1RN* DNAm edited iMacs, including increased levels of genes related to cell division (*BRCA1* and *CDK1*) and metabolism (*HK2* and *PGK1*), as well as reduced levels of B-(*LEF1* and *CD79A*) and myeloid-(*ITGAM* and *CD14*) related genes **(Extended Data Fig. 3f)**. Furthermore, the shIL1RN iMacs exhibited as well reduced levels of the myeloid surface markers CD11b and CD14 and enhanced phagocytosis capacity compared with control iMacs **(Fig. 3g-h and Extended Data Fig. 3g)**.

Utilizing an orthogonal approach, we have confirmed the findings regarding the involvement of *IL1RN* and perhaps, by extension, of the IL1 pathway in the establishment of the C/EBPα-driven myeloid cell fate and acquisition of enhanced phagocytic capacity. Thus, this suggests that the IL1 pathway may be necessary for the proper acquisition of myeloid cell fate during human hematopoiesis.

### C/EBPα may activate key IL1 pathway-related GREs during the acquisition of the myeloid cell fate

Based on our observation of IL1RN’s dependency on the correct acquisition of the myeloid cell fate in our cellular model, we analyzed its epigenomic regulation during the process to gain insight into its transcriptional control. We observed that C/EBPα, which drives the transdifferentiation process, binds to the *IL1RN* promoter as early as 24 hours. This binding occurs before the DNA demethylation of the region and the transcriptional activation **(Fig. 1g and Fig. 2k)**, but it temporally coincides with a significant epigenome rewiring, as indicated by increased chromatin accessibility and activation (detected by ATAC-seq and H3K27ac decoration, respectively) at the C/EBPα-bound region **(Fig. 3i upper panel).**

Another IL1 pathway-related GRE at *IL1B* locus was found to undergo demethylation during the transdifferentiation process **(Fig. 1e)**. This GRE is a proximal enhancer (PE) of the gene (PE2 at -5.5kb from the TSS) that is bound by C/EBPα along with its promoter, and yet another proximal enhancer (PE1 at -3kb from the TSS) as early as 24h of transdifferentiation (**Extended Data Fig. 3h upper left panel)**. In B cells, both the *IL1B* promoter and the PE1 are already accessible and devoid of DNAm **(Extended Data Fig. 3h, upper right panel)**. However, the binding of C/EBPα to the PE2 occurs in a closed chromatin region that only becomes accessible after its binding, coinciding with the loss of DNAm in the region **(Extended Data Fig. 3h, upper right panel)**. The C/EBPα binding to the *IL1B* promoter, PE1 and PE2 regions might finally favor the establishment of a temporary super-enhancer, which is observed to cover the PEs and the entire gene body (approximately 10kb). This is indicated by the coverage of the whole region with H3K4me1 and H3K27ac, ultimately leading to increased transcriptional output, as shown by the enhanced activation of the promoter (by H3K4me3 deposition) **(Extended Data Fig. 3h upper left panel)**.

The observed DNA demethylation and chromatin activation events at the *IL1RN* and *IL1B* genes upon C/EBPα binding to their GREs strongly suggest that this transcription factor plays an important role in controlling the IL1 pathway activity during the establishment of the myeloid cell fate. To gain insight into the physiological process, we studied the transition from mouse hematopoietic stem progenitor cells (HSPCs) to granulocyte-macrophage progenitors (GMPs) driven by C/EBPα^37^. We only observed chromatin activation at the *Ilrn* and *Il1b* loci at the GMP stage, specifically at the C/EBPα-bound regions, coinciding with the GREs described in the human cellular cell fate model **(Fig. 3i and Extended Data Fig. 3h, lower panels)**.

Altogether, these analyses reveal an unidentified mechanistic role for C/EBPα as a potential crucial transcriptional regulator of the IL1 pathway during both human and murine myeloid cell fate determination.

### *IL1RN* promoter DNAm editing enhances the cellular transcriptional response to the IL1 pathway activation

Since IL1RN acts as a negative regulator of the IL1-signaling pathway^38^, the *IL1RN* DNAm edited (sgIL1RN) iMacs that are generated by the end of the transdifferentiation process might have this pathway hyperactivated in the steady state. We observed an increased nuclear localization of the NFKB subunit p65 in sgIL1RN iMacs, indicating increased transcriptional activity of the IL1-signaling pathway in the steady state **(Extended Data Fig. 4a)**. Based on these findings, we questioned whether sgIL1RN iMacs may exhibit increased sensitivity to inflammatory stimuli and respond differently to this challenge compared to CTRL iMacs. To test this hypothesis, dCas9-DNMT3A CTRL and sgILRN iMacs were treated with the IL1-pathway agonist, IL1β, for different time points and then collected for further analyses **(Fig. 4a)**. Upon IL1β stimulation, the nuclear localization of p65 is further enhanced in the sgIL1RN compared to CTRL iMacs **(Fig. 4b)**, validating the notion of a hypersensitive state for the activation of the IL1-pathway. To further characterize it, we assessed the transcriptomic profiles of dCas9-DNMT3A CTRL and sgILRN iMacs untreated (0h) or treated with IL1β for 3 and 16 hours. The original differences in gene expression, which were already noticeable in the unstimulated cells (0h), persisted when the two types of iMacs were exposed to IL1β for a short period (3h) **(Fig. 4c and Extended Data Fig. 4c-d)**. However, these differences disappeared after longer IL1β exposure (16h) **(Fig. 4c and Extended Data Fig. 4c-d).** This may be because the prolonged stimulation of the IL1 pathway resulted in the activation of feedback inhibitory loops previously described to operate in this pathway for resolving inflammation^39^. To understand how the DNAm editing of *IL1RN* impacts the activation of important transcriptional regulatory networks (regulons), we used DoRothEA (Discriminant Regulon Expression Analysis). DoRothEA is a manually curated human regulon that estimates the activities of transcription factors in individual samples based on the expression of their target genes^40^. The analysis showed higher activity in regulons related to important myeloid (SPI1 or RUNX1) and inflammatory (RELA) transcription factors and lower activity in regulons associated with the interferon pathway (STAT2 or IRF9) in sgIL1RN compared to CTRL iMacs, both at 0h and mostly at 3h after I IL1β treatment **(Fig. 4d and Extended Data Fig. 4e)**. At 16h of IL1β, the transcriptional activity detected with DoRothEA mirrored the patterns described at 0 and 3h **(Extended Data Fig. 4e)**. This is likely due to the previously described IL1-feedback inhibitory loops^41^.

**Figure 4.**
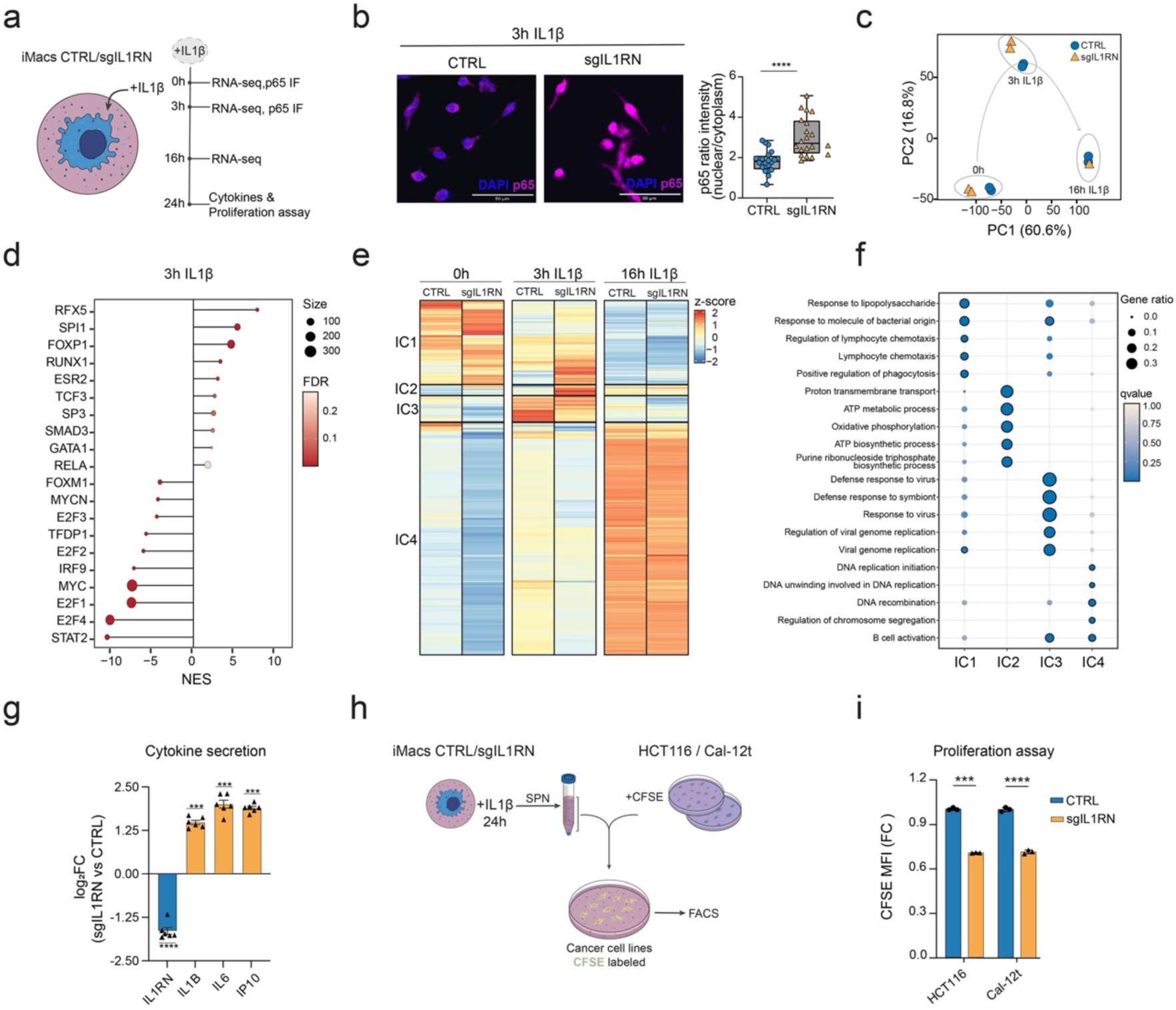
*IL1RN* DNAm editing increases cell’s sensitivity to inflammatory stimuli. **(a)** Schematic of the IL1β treatment experiment. **(b) Left:** Representative IF showing p65 (red) and DAPI (blue) signals in sgIL1RN and CTRL iMacs treated with IL1β for 3h. **Right:** quantification of p65 nuclear versus cytoplasmic localization signal in sgIL1RN and CTRL iMacs treated with IL1β for 3h. Unpaired two-tailed Student’s t-test, n=20 cells per group, mean s.e.m., (****p< 0.0001). **(c)** Principal Component Analysis (PCA) displaying the transcriptomic dynamics of sgIL1RN and CTRL iMacs treated with IL1β. **(d)** Lollipop plot depicting the TF activity predicted from mRNA expression of target genes with DoRothEA v2.0^40^ in sgIL1RN and CTRL iMacs treated with IL1β for 3h. Top 10 most significant (FDR < 0.05) transcriptional target genes from positive and negative normalized enrichment are shown. Lollipop size indicates the total number of DEGs regulated by each transcriptional regulon. **(e)** Clustering of the transcriptional dynamics (by RNA-seq) observed during the IL1β-treatment in sgIL1RN and CTRL iMacs. Average scaled normalized counts of differentially expressed genes (padj < 0.05) are represented at each timepoint (n=2 biologically independent replicates). Black lines show splitting by k-means clustering. IL1β-treatment Clusters (IC1-4). **(f)** Balloon plot depicting the Gene Ontology (GO) enrichment analysis for the genes associated with clusters in e. The top 5 most significantly over-represented biological processes (BP) terms for each cluster are plotted. Ratio of genes of interest over all unique genes (GeneRatio) and qvalue are shown. Significant terms (qvalue < 0.05) are highlighted with a black stroke. **(g)** Normalized levels of secreted cytokines in sgIL1RN iMacs treated for 24h with IL1β. One-way ANOVA with Dunnett’s post-hoc correction, n=6, mean s.e.m., (***p < 0.001; ****p < 0.0001). **(h)** Schematic of the experimental setup used to assess the impact of the cytokines produced by the sgIL1RN iMacs in cancer cell lines’ proliferation (measured by CFSE retention). **(i)** Plot showing the normalized Mean Fluorescence Intensity (MFI) for the CFSE signal (4 days after staining) in cancer cell lines exposed to the supernatant of sgIL1RN or the supernatant of CTRL iMacs. Unpaired two-tailed Student’s t-test, n=20 cells per group, mean s.e.m., (***p < 0.001; ****p< 0.0001).

To investigate the impact of *IL1RN* DNAm editing on the transcriptomic kinetics, all the genes differentially expressed between sgILRN and CTRL iMacs throughout the IL1β-treatment were selected and clusterized into 4 major groups (IL1βI-Clusters -IC- 1 to 4) **(Fig. 4e)**. While genes in the IC1 cluster are downregulated, the genes in the remaining clusters are either transiently upregulated (IC3) or continuously upregulated (IC2 and IC4) along the treatment **(Fig. 4e and Extended Data Fig. 4f)**. *IL1RN* DNAm editing led to impaired downregulation kinetics in the genes contained in IC1, as exemplified by the myeloid gene *CD14* **(Fig. 4e and Extended Data Fig. 4f)**. Of note, IC1 is enriched in genes related to “Response to lipopolysaccharide” (LPS), “Lymphocyte chemotaxis”, or “Regulation of phagocytosis” among others **(Fig. 4f)**. In IC2, *IL1RN* DNAm editing induced a strong and transient gene upregulation at 3h of IL1β -treatment (instead of the continuous upregulation observed in CTRL cells) **(Fig. 4e and Extended Data Fig. 4f)**. Genes within this cluster are related to metabolic biological processes (including “ATP metabolic process” or “Oxidative phosphorylation” among others), such as the mitochondrially encoded ATP synthase 6, *MT-ATP6* **(Fig. 4f and Extended Data Fig. 4f)**. Genes in IC3 showed lower levels at 0h in *IL1RN* edited cells, and the editing also hindered their full activation at 3h **(Fig. 4e and Extended Data Fig. 4f)**. These genes are related to viral response biological processes (including “Defense response to virus” or “Viral genome replication”), as exemplified by the kinetics of the interferon-induced genes 2’-5’-oligoadenylate synthetase 2 and 3 (*OAS2* and *OAS3*) **(Fig. 4f and Extended Data Fig. 4f)**. Finally, IC4-included genes are characterized by lower basal levels of expression (at 0h) in *IL1RN* DNAm edited cells and progressive recovery to reach the levels displayed in CTRL cells upon IL1β treatment **(Fig. 4e and Extended Data Fig. 4f)**. IC4 genes are related to cell division biological processes (including “DNA replication initiation” or “DNA recombination”), as exemplified by the expression kinetics of the essential cell cycle regulators *CDC45* and *CDC6* **(Fig. 4f and Extended Data Fig. 4f)**.

Altogether, the transcriptional analysis performed suggests that by modifying the DNAm status of the *IL1RN* promoter, we altered the cellular response to the IL1β-mediated inflammatory stimulus.

### *IL1RN* promoter DNAm editing alters critical cellular inflammatory properties

In line with the observed altered transcriptional response to the IL1 pathway activation, the sgIL1RN iMacs displayed a distinct cytokine secretory profile after a 24-hour IL1β-treatment **(Fig. 4g and Extended Data Fig. 4g)**. This profile was characterized by higher levels of well-known proinflammatory cytokines, including IL1β, IL6, and IP10 **(Fig. 4G)**. All of them showed an average increase of at least 2-fold in their levels, with the highest differential accumulation observed for IL6 (>3.5-fold) **(Fig. 4g)**. The impact of the DNAm editing on *IL1RN* was also detectable at the cytokine level. Even after 24 hours of IL1β-treatment, when feedback loops are operating, edited cells displayed >2-fold decrease in secreted IL1RN levels **(Fig. 4g)**.

Next, we explored the intriguing possibility that modifying the DNAm status of a single gene promoter (*IL1RN*) might be sufficient to modulate cancer cell growth. This is because macrophages have been shown to potentially regulate cancer development in the tumor microenvironment by producing different sets of cytokines^42^. To that end, we collected the supernatant of CTRL and sgIL1RN iMac after 24-hour IL1β-treatment **(Fig. 4h)**. Then, we labeled cells from different cancer origins (HCT116: colon cancer and Cal-12t: non-small cell lung carcinoma) with the proliferation dye Carboxyfluorescein Succinimidyl Ester (CFSE) and cultured them using the supernatant previously collected from the CTRL and sgIL1RN iMacs **(Fig. 4h)**. We observed that cells from both cancer origins when grown in the IL-1β, IL-6, IP-10-rich supernatant of the sgIL1RN iMacs, retain an average of 30% less of the CFSE signal 4 days after staining **(Fig. 4i)**. This means that the differential cytokine profile secreted by the *IL1RN* DNAm edited cells favors the proliferation of these cancer cells. Of note, macrophages within the bone marrow have recently been described as potential drivers of leukemic transformation^43^.

In summary, these analyses suggest that modifying the DNAm levels of the *IL1RN* promoter has led to the production of macrophages with a pro-inflammatory cytokine release profile that may have a distinct potential to support cancer cell growth.

## DISCUSSION

The link between DNAm and gene expression has been widely studied, with many reports showing a negative correlation between DNAm levels and gene expression, especially at promoter sites^44^. However, reports describing causal relationships between DNAm, expression, and cellular phenotypes are scarce^30^.

Here, we have utilized the highly efficient and homogenous reprogramming of B cells into macrophages to conduct an integrative analysis of DNAm and transcriptional dynamics. Our findings revealed correlative changes, specifically in gene regulatory regions experiencing a loss of DNAm. These changes were associated with genes related to myeloid cell fate and inflammation, such as the promoter of the Interleukin-1 receptor antagonist (*IL1RN*) gene. Then, we targeted this region with CRISPR/dCas9 DNAm editing tools and found that altering the methylation status of the *IL1RN* promoter region was sufficient to trigger significant changes in gene expression, leading to distinct cell fate outcomes and responses to inflammatory signals.

Analyzing the dynamics of DNAm, we observed that the changing regions were unevenly distributed along the different chromosomes, with the largest changes occurring on ChrX **(Extended Data Fig. 1b)**. Notably, about 32% of the DNAm signal on ChrX is rewired during this process, which is 2.3 times higher than the genomic average changes. Specifically, In this chromosome, 2 out of 3 regions change their DNAm by losing signal in the induced macrophage state. As our B cells are derived from the RCH-ACV female B-ALL cell line^45^, the significant changes observed in ChrX may indicate either a partial reactivation of the inactive ChrX during the cell fate conversion or abnormal hypermethylation of the active ChrX in B cells. On the one hand, it has been described that immune cells may not maintain the inactivation of ChrX as efficiently as other somatic cells^46–48^. This could lead to the overactivation of immune-related genes abundantly located on this chromosome, ultimately contributing to the development of autoimmune diseases that have a higher incidence in female individuals^49^. On the other hand, hypermethylation of ChrX and reduced levels of the master regulator of ChrX dynamics, Xist, have been described as mechanisms that promote progression in breast and blood cancers^50,51^. Therefore, the loss of DNAm at ChrX may contribute to the loss of tumorigenicity associated with the B-to-macrophage transdifferentiation process.

In addition, during the transdifferentiation process, we observed a loss of DNAm at GREs associated with myeloid genes. These GREs included key transcription factors such as *KLF4* or *MAFB*, chromatin regulators such as the DNA dioxygenase *TET2*, and genes associated with inflammation such as the interferon-gamma receptor gene *IFNGR1* or the cytokine-coding genes *IL1B*, *IL1RN* and *IL6* **(Fig. 1e)**. While the involvement of KLF4^52^, MAFB^53^ and TET2^12,54,55^, in immune-related cell fate decisions has been documented previously, the role of the inflammatory pathways has not been extensively explored. The IL6 pathway was found crucial for establishing proper pluripotency during the conversion of mouse embryonic fibroblasts^56,57^, or murine pre-B cells^58,59^, into induced pluripotent stem cells (iPSCs). However, it was found to be dispensable for myeloid acquisition during the conversion of murine pre-B cells into induced macrophages^59,60^. Conversely, the IL1 pathway has not been studied in the context of experimentally induced cell fate conversions. Here, we identified GREs associated with the IL1 pathway-related genes *IL1B* and *IL1RN* losing DNAm during human B-to-macrophage transdifferentiation **(Fig. 1e).** These genes encode for the agonist and the antagonist of the pathway, respectively^61^. Interestingly, these IL1 pathway-related GREs losing DNAm coincide with C/EBPα binding sites. Notably, its binding potentially triggers their DNA demethylation and associated chromatin activation, finally leading to their increased transcriptional activity (**Fig 3i and Extended Data Fig. 3h)**. The observed DNA demethylation and chromatin activation by C/EBPα at these GREs might be mediated by its ability to act as a pioneer transcription factor engaging a broad range of chromatin modifiers^52,62^, including TET2^12^.

The IL1-related GREs that C/EBPα binds to and activates in the human conversion system are also occupied by this factor during the homeostatic differentiation of murine HSPCs to GMPs **(Fig 3i and Extended Data Fig. 3h)**. This suggests that C/EBPα could be a significant transcriptional regulator of the IL-1 pathway during physiological myeloid establishment. By reducing the levels of *IL1RN* using DNAm editing and shRNA techniques, we found that in our cellular model of human myeloid cell fate acquisition, there was an abnormal shift in the myeloid cell fate and enhanced phagocytic capacity **(Fig. 3a-h and Extended Fig. 3a-g)**. All of these changes are consistent with the imbalanced production of blood cells observed in mice that were chronically exposed to IL-1β^35^ or those with an *Il1rn* deficiency^36^. This is also in line with the observation that lower levels of *IL1RN* are associated with reduced survival rates in AML patients, particularly for the more differentiated subtypes M4-M5^36^. Based on the strong association we have proven between the expression of *IL1B* and *IL1RN* and the methylation status of some of their GREs, it is likely that the abnormal activation of the pathway observed in AML patients is caused by an abnormal methylation of GREs associated with the *IL1RN* or *IL1B* genes.

In mature myeloid cells, the IL1 pathway plays a crucial role in responding to inflammatory stimuli^29^. Overactivity in this pathway leads to various immune disorders that can be managed by treating patients with a synthetic form of IL1RN known as anakinra^63,64^. One such immune disorder is the Deficiency of the Interleukin-1-Receptor Antagonist (DIRA), a rare but severe autoinflammatory disease characterized by mutations in the *IL1RN* gene^65^. However, in most IL1-related autoinflammatory conditions, such as cryopyrin-associated periodic syndromes (CAPS), there are no mutations in the *IL1RN* or *IL1B* genes. Rather, hyperactivation of the pathway in CAPS may be caused by abnormal methylation at their gene promoters. This molecular condition has been shown to be correctable by anti-IL-1β treatment^66^. Thus, it suggests that inflammatory responses involved in certain pathological conditions might be fine-tuned through the methylation status of the *IL1RN* gene. We have proven this hypothesis by using a DNAm editing tool to induce hypermethylation of the *IL1RN* promoter in a physiologically relevant macrophage cellular model^67,68,26^. The DNAm editing of the *IL1RN* promoter results in a basal proinflammatory state, which causes cells to be hyperresponsive to IL1β stimulation **(Fig 4b and Extended Fig. 4a)**. This leads to an abnormal transcriptional response to the stimulus, varying secretion of proinflammatory cytokines, and different ability to support cancer cell proliferation **(Fig 4c-i and Extended Fig. 4c-g)**.

While major drug agencies have already approved a CRISPR/Cas9-based DNA editing treatment^67^, therapies for editing the epigenome are being delayed due to initial off-target effects and challenges in accurate delivery into the targeted cells^68^. Despite these limitations, preclinical studies show promise in using DNAm editors, for instance, in reversing disease-associated abnormal DNAm events in neurological conditions^32,69^. Notably, a recent breakthrough in epigenome therapy achieved permanent silencing of a gene involved in cholesterol homeostasis directly in the mouse liver^70^. Therefore, based on recent technological developments, an epigenome editing therapy to modulate the DNAm levels of the *IL1RN* promoter in human myeloid cells might be a reality in the mid-term. Based on our findings, such therapy might be instrumental in fine-tuning the activity of the IL1 pathway in a broad range of human conditions, from controlling autoinflammatory processes to modulating the growth of leukemia or solid cancers.

## METHODS

### Cell lines and cell culture

BLaER cells are derived from a human B-cell precursor leukemia cell line (RCH-ACV) that stably expresses the myeloid transcription factor C/EBPα fused with the estrogen receptor (ER) and labeled with GFP^24^. BlaER cells and subclones were grown in suspension in RPMI 1640 (GIBCO) supplemented with 10% heat-inactivated FBS (GIBCO), 1X Penicillin-Streptomycin (GIBCO), 1X L-glutamine (GIBCO) and 0,1X β-mercaptoethanol (GIBCO). Culture medium was replaced every 2-3 days upon counting in a hemocytometer and using trypan blue exclusion dye to discriminate between live and dead cells. Then, cells were seeded at 2 x10^5^ cells per ml into an appropriate tissue culture flask.

HEK-293T cells were grown in Dulbecco’s Modified Eagle’s Medium (DMEM) (+) D-glucose supplemented with 10% heat-inactivated FBS, 1X L-glutamine, and 1X Penicillin-Streptomycin.

For optimal growth, all cell lines were kept in a 5% CO2 humidified atmosphere at 37 °C.

Cells were checked for mycoplasma infection every month and tested negative.

### Transdifferentiation of human B cells into macrophages

To induce transdifferentiation of human leukemic B cells (B cells) into induced macrophages (iMacs), 2 x10^5^ BLaER cells were seeded in a 12-well plate. To activate C/EBPα, 100 nM 17-beta estradiol (E2) (Sigma Aldrich), human IL-3 (10 ng per ml) (PeproTech) and human M-CSF (10 ng per ml) (PeproTech) were added to the medium to favor the conversion.

### Lentiviral production and cellular transduction with the DNA methylation editing tools

To transduce BlaER cells with the DNA methylation editing tools, low-passaged HEK293T cells were seeded for transfection. Lentiviruses were produced in these cells by co-transfecting them with lentiviral plasmids VSV-G, psPAX2, and a transfer vector (Fuw-dCas9-DNMT3A-P2A-tagBFP -Addgene #84569-; Fuw-dCas9-TET1-P2A-tagBFP -Addgene #108245- and pLV GG hUBC-dsRED -Addgene #85034-) using calcium phosphate. Supernatants containing lentiviral particles were collected and filtered at 48- and 96-hours post-transfection and concentrated by centrifugation at 70.000 x g for 2 h at 10 °C. BLaER cells were then spin-infected (1000 x g; 32 °C for 90 min) with the concentrated viruses. Single-cell sorting of the BFP+ population was performed to generate stable dCas9-DNMT3A-P2A-BFP (hereafter dCas9-DNMT3A) or dCas9-TET1-P2A-BFP (hereafter dCas9-TET1) clones. Single sorted cells were then grown at 37°C in 20% FBS-RPMI medium for 14-20 days to generate individual clones containing the dCas9-DNMT3A or the dCas9-TET1 epigenome editing tool. To modify the methylation status, dCas9-DNMT3A and dCas9-TET1 clones were selected (based on their dCas9/BFP expression levels). Next, selected clones were infected with lentiviruses harboring simultaneously 4 sgRNAs targeting the *IL1RN* gene promoter (sgIL1RN) and scramble regions (sgCTRL).

The design of the single guide RNAs (sgRNAs) targeting the *IL1RN* promoter (chr2:113,127,440-113,127,701) and CTRL regions was performed using Benchling sgRNAs design tool for CRISPR (https://www.benchling.com/crispr). sgRNAs close to a protospacer adjacent motif with 5’-NGG-3’ were selected with the best on-target and off-target scores. Multiplex sgRNA-dsRED (pVL GG hUBC-dsRED plasmid -Addgene # 84034-) construct was cloned using a two-step protocol as described by Engler and coworkers^73,74^. Briefly, each protospacer was first annealed and cloned into the desired expression vector [phH1-gRNA (Addgene # 53186); ph7SK-gRNA (Addgene #53189); pmU6-gRNA (Addgene # 53187), and phU6-gRNA (Addgene #53188)] using a BbsI restriction enzyme site. Next, four promoter-gRNA cassettes were cloned into the lentiviral destination vector GG hUBC-dsRED using Golden Gate assembly^75^. Plasmids were sequenced using the M13 Reverse primer. Single gRNA sequences are listed in **Table 1**.

### CSFE proliferation assay in cancer cells

To assess cell proliferation, dCas9-DNMT3A CTRL and sgIL1RN iMacs were treated for 24h with IL1β (1ng per ml). Then, 2 million HCT-116 or Cal-12t cancer cells were stained with CFSE (Invitrogen Cat #C34570) following the manufacturer’s instructions. Next, cancer cells were seeded at 10,000 cells per ml and grown with supernatant collected from CTRL or sgIL1RN iMacs **(Fig. 4h)**. Cells were analyzed for CFSE levels at the FACS Canto II flow cytometer at days 0 and 4. Data was analyzed with FlowJo software.

### Immunocytochemistry and microscopy for p65

To assess p65 cellular localization, dCas9-DNMT3A CTRL and sgIL1RN iMacs were seeded at 0,8 x 10^6^ cells per ml on poly-lysine coated coverslips. After 2 days, iMacs were treated with IL1β (1ng per ml) for 3h. Next, cells were fixed with 4% paraformaldehyde (in PBS) for 20 min and permeabilized with PBS + Triton X-100 0.5% for 10 min. Coverslips were washed twice with PBS, blocked with blocking solution (PBS + BSA 4% + 0.025% Tween 20) and incubated with an anti-p65 antibody (1:200 Abcam Cat #ab16502) overnight at 4°C. After washing, cells were incubated with an anti-rabbit Alexa Fluor 647 (1:300 Invitrogen Cat #A21245) for 1 h at room temperature. After four washes with PBS, cells were stained with DAPI 2 μg per ml and mounted into slides using Vectashield (Vector Laboratories Cat # H-5700-60). Images were obtained with a Leica TCS-SL confocal microscope. ImageJ was used to quantify the mean intensity of the nuclei/cytoplasm ratio for p65 signal. 20 cells were counted for each sample.

### shRNA-mediated IL1RN silencing

For constitutive shRNA-mediated gene silencing, oligonucleotide pairs encoding *IL1RN* shRNAs were annealed and cloned into pSICOR-PGK-puro (Addgene #12084). pSicoOligomaker 1.5 (https://venturalaboratory.com) was used to select and design oligos. Knockdown was confirmed by qRT-PCR. The following shRNAs were used: Luciferase shRNA (shLuc): CCTAAGGTTAAGTCGCCCTCG IL1RN shRNA_1 (sh1): GCGTCATGGTCACCAAATT IL1RN shRNA_2 (sh2): GTACTATGTTAGCCCCATA BlaER cells were transduced with shLuc or shIL1RN lentiviruses and selected with puromycin (1 µg per ml) for 2 days.

### Phagocytosis assay

Phagocytosis assays were performed using dCas9-DNMT3A CTRL and sgIL1RN iMacs. The assays were also performed in iMacs containing shRNA (shLuc and shIL1RN_1/2). In both cases, cells were seeded at a concentration of 0.5 x 10^6 cells per ml in DMEM media. Next, Fluoresbrite carboxyl bright blue beads (1 µm, Polysciences Cat # 17458-10) were added to the media (300 beads per cell) and incubated for 24h before FACS analysis. Data was analyzed with FlowJo software.

### Surface markers profiling

Stainings for CD14 and CD11b were conducted in dCas9-DNMT3A CTRL and sgIL1RN iMacs, as well as in iMacs with shRNAs (shLuc and shIL1RN_1/2). Briefly cells were collected and incubated with Human FcR Binding Inhibitor (eBiosciences Cat #16916173) to avoid unspecific staining. Next, cells were stained against the macrophage marker CD11b (1:200 of CD11b-APC BD Pharmingen Cat #550019) and CD14 (1:200 of CD14-APC BD Pharmigen Cat #561383) and resuspended in PBS containing DAPI as a viability marker.

### Western Blotting

Cells were lysed in Laemmli sample buffer (8% SDS, 40% glycine, 20% β-mercaptoethanol, 0,250 M Tris pH 6.8, 0,008% bromophenol blue) and boiled for 5 min at 95°C. Protein extracts were separated by electrophoresis and transferred to a nitrocellulose membrane. Membranes were blocked with 5% non-fat milk or 5% BSA for 1 hour at RT. Then, membranes were incubated overnight at 4°C (while shaking) with the following primary antibodies: goat anti-IL1RN (RD Systems Cat # AF280NA) in 1:1000 in blocking solution and mouse anti-β actin (MERCK Cat # A1978) and rabbit anti-vinculin (Cell Signaling Cat #18799) 1:5000 in blocking solution. Membranes were washed 3 times in TBS-Tween before incubation with a secondary antibody anti-mouse IgG Alexa Fluor 790 nm (Invitrogen Cat #A11375), anti-goat IgG Alexa Fluor 800 nm (Invitrogen Cat# A32930) or anti-rabbit IgG Alexa Fluor 680 nm (Invitrogen Cat # A-21109) 1:5000 in blocking solution for 1 hour at RT. Finally, membranes were developed in an Odyssey CLx system. Image Studio Lite 5.2 (Li-COR) was used for visualization and quantification analysis.

### Cytokine measurements

The LEGENDplex™ Human M1/M2 Macrophage Panel (10-plex) (BioLegend Cat# 740508) was used to quantify proinflammatory (IL-12p70, TNF-α, IL-6, IL-1β, IL-12p40, IL-23 and IP-10) and anti-inflammatory (IL-10, TARC, and IL-1RA) cytokines in the supernatant of dCas9-DNMT3A CTRL and sgIL1RN iMacs treated with IL1β for 24h. The recommended filter plate method was used, and all steps were followed per the manufacturer’s protocol. Samples were acquired in a FACS Canto II flow cytometer, and LEGENDplex™ Data Analysis Software (BioLegend) was used for data analysis according to the manufacturer’s recommendations. Samples and standards were run in duplicate.

### Real-time quantitative Polymerase Chain Reaction (RT-qPCR)

Total RNA was extracted using Trizol (eBioscences). 500 ng of total RNA were converted into cDNA using the RNA to cDNA kit (Applied Biosystems) following the manufacturer’s instructions. Real-time quantitative PCR reactions were performed using SYBR Green reagent and analyzed using QuantStudio 5 System (Applied Biosystems). *HPRT and B2M* were used as housekeeping genes. Unpaired student’s t-test was used to determine statistical differences in gene expression among the different samples tested (t-test, ***p<0.001). Normality and homogeneity in variance were assumed for RT-qPCR experiments with biological triplicates. Primer sequences are listed in **Table 2**.

### RNA-seq

For the transdifferentiation experiment, approximately 1–2 × 10^6^ of dCas9-DNMT3A CTRL or sgIL1RN cells were collected in duplicates at 0, 3 and 7 days after C/EBPα induction. For the IL1β treatment experiment, dCas9-DNMT3A CTRL or sgIL1RN iMacs were collected in duplicates at 0, 3 and 16h after IL1β (1 ng per ml) treatment. Total RNA was extracted with the RNeasy Mini Kit (QIAGEN Cat#74104) according to manufacturer’s instructions and quantified with NanoDrop spectrophotometer. 0.2-0.5ug of RNA was used for mRNA sequencing with poly-A enrichment. Briefly, quality control with Fragment analyzer (Agilent) was performed before library preparation to ensure proper RNA integrity (RIN>7). A DNABSEQ Eukaryotic Strand-specific mRNA library protocol was used for library preparation. Then, libraries were sequenced in a DNABSEQ-G400 sequencer using a pair-end 150-bp protocol. More than 35 million reads were obtained for each sequenced sample.

### Bisulfite conversion (BS) and pyrosequencing

Sample DNA methylation status was assessed by bisulfite pyrosequencing. Briefly, 1 ug of genomic DNA was bisulfite (BS)-converted using the EZ DNA Methylation Gold Kit (Zymo Research Cat# D5006) following the manufacturer’s instructions. BS-treated DNA was PCR-amplified using the IMMOLASE DNA polymerase Kit (Bioline). Primers used for the PCR were designed with PyroMark Assay Design 2.0 software (QIAGEN) and listed in **Table 3**. PCR products were pyrosequenced with the Pyromark Q48 system (QIAGEN), according to the manufacturer’s protocol and analyzed with PyroMark Q48 Autoprep (QIAGEN).

### Whole Genome Bisulfite-seq

Genomic DNA was extracted from 1 million cells at 0, 24, 96, and 168 hours of transdifferentiation using the DNeasy Blood & Tissue kit (QIAGEN Cat #69504) following the manufacturer’s instructions and quantified using Qubit dsDNA (Invitrogen Cat #Q32851). Cytosine conversion, library preparation and sequencing were done by the provider of the sequencing services. Briefly, genomic DNA was fragmented to 200-400 bp and bisulfite treated. For library construction, sequencing adapters were ligated, followed by double-strand DNA synthesis and PCR amplification. Next, libraries were sequenced on Illumina HiSeqTM2500 using a pair-end 150-bp protocol rendering >70Gb/sample. Raw data quality assessment was performed, and low-quality reads were trimmed.

### DNA methylation arrays

To assess potential off-target DNA methylation events caused by the dCas9-DNMT3A tool, we utilized the Infinium MethylationEPIC v2.0 Bead-Chip arrays (Illumina Cat #20087706). This platform allows the interrogation of around 935,000 CpG sites per sample at single-nucleotide resolution, covering 99% of the reference sequence (RefSeq) genes. 1 million CTRL or sgIL1RN cells were collected at 0 and 3 days after C/EBPα induction, with 4 biological replicates for each group. Genomic DNA was extracted and bisulfite-converted as previously described and used to hybridize the methylation arrays following the manufacturer’s instructions. Raw files (IDAT files) were provided by the Genomics Unit of the Josep Carreras Research Institute (Barcelona).

### Chromatin immunoprecipitation (ChIP)

ChIP experiments were performed as previously described^12^. Briefly, 30 million cells were cross-linked with 1% formaldehyde (Sigma) in RPMI medium while rotating for 10 minutes at room temperature. To stop fixation, glycine was added to a final concentration of 0,125M and rotated for 5 minutes. Next, collected cells were washed twice with ice-cold PBS and resuspended in cold IP buffer (1 volume SDS buffer (100mM NaCl, 50mM pH8.1 Tris-HCl, 5mM pH8 EDTA, 0.2% NaN3, 0.5% SDS): 0.5 volume Triton dilution buffer (100mM NaCl, 100mM pH8.6 Tris-HCl, 5mM pH8 EDTA, 0.2% NaN3, 5% Triton X-100) supplemented with proteinase inhibitors (Roche Cat #118733580001). Chromatin was sheared to 100-300bp fragments on a Bioruptor pico sonicator (Diagenode) at 4°C for 13 cycles with 30 sec on and 30 sec rest in 15 ml Bioruptor Tubes (Diagenode Cat #C30010017) with 650 mg of beads (Diagenode Cat #C03070001). After sonication, cell lysate was spun down at 20.000xg for 20 min at 4°C to remove debris and 5% of supernatant was saved as input. Chromatin was conjugated with 10ug of anti-Cas9 antibody (Active Motif Cat #61757) while rotating ON at 4°C. The next day, 50 ul of Dynabead A/G mix were blocked for 2h at 4°C with BSA 5mg per ml while rotating and added into antibody-cell lysate mixture for immunoprecipitation for 3h at 4°C in rotation. Chromatin-antibody-bead complexes were washed three times with ice-cold low salt buffer (50mM pH7.5 HEPES, 140mM NaCl, 1% Trito X-100) and once with ice-cold high salt buffer (50mM pH7.5 HEPES, 500mM NaCl, 1% Triton X-100). Bound protein-DNA complexes were de-crosslinked in elution buffer (1% SDS, 0.1M NaHCO3) by overnight incubation at 65°C with shaking at 1300 rpm. The next day, the eluted portion was treated with RNAse for 1h at 37°C and then with Protein K for 2h at 65°C. Finally, DNA was purified by phenol:chloroform:isoamyl alcohol (25:24:1) extraction and ethanol precipitated.

For ChIP-qPCR analysis, DNA was diluted 1:10, and relative enrichment was calculated as a percentage of input with the following formula (100*2^(Adjusted input - Ct (IP)). Oligonucleotide sequences are indicated in **Table 4**.

For ChIP-seq, samples were quantified with Agilent 2100 before library preparation. Library preparation and sequencing were performed by the sequencing service provider using a DNABSEQ-G400 sequencer and a SE50 protocol.

### Bioinformatic analyses

All sequencing data obtained were mapped onto the human genome assembly hg38 (Ensembl GRCh38) and analyzed with R (4.2.1) using packages from the Bioconductor suite (v3.0)^76^. For peak calling, regions overlapping the ‘Encode blacklist’ regions were removed^77^, as well as mitochondrial reads. Peaks were annotated to genomic features in R with the package ChIPseeker (v1.32.1)^78^, using Benjamini-Hochberg (BH) FDR corrections. All GO enrichment analyses were performed using the clusterProfiler package (v4.4.4)^79^. Bigwig tracks were generated using DeepTools BamCoverage (3.3.1)^80^. Motif enrichment analyses were performed using the function findMotifsGenome.pl from the package HOMER (v0.2)^81^. The integration of DNA methylation and ATAC-seq data was carried out using the bedtools (v2.31.1)^82^, package. Heatmaps and clustering analyses were performed using the ComplexHeatmap (v2.10.0) package^83^. The Multidimensional Scaling (MDS) and Principal Component Analysis (PCA) were performed using the limma (v3.52.2)^84^, and factoextra packages (v1.0.7), respectively. Upset plots were generated using the upsetR package (v1.4.0)^85^. The correlation analyses were performed and plotted with the R package corrplot (v0.92). The RNA-seq MA plots were generated using the R package DESeq2 (v1.36.0)^86^. The remaining plots were generated using the R package ggplot2 (v3.4.2). Genome-wide statistical analyses were performed using a two-tailed student t-test to compare two groups using the R package stats (v4.2.1).

### Bisulfite-seq analysis

Raw sequence reads from WBGS libraries were trimmed to remove poor-quality reads and adapter contamination using the package Trimmomatic (v0.36)^87^. The remaining sequences were mapped using Bismark (v0.16.3)^88^, to the human reference genome GRCh38 in paired-end mode. Reads were then deduplicated, and CpG methylation calls were extracted from the deduplicated mapping output using the Bismark methylation extractor in paired-end mode. The estimation of DNA methylation levels was calculated by dividing the count of reads identifying a cytosine (C) by the sum of reads identifying either a cytosine (C) or a thymine (T). Only CpGs with coverage equal to or higher than 5X in all tested samples were used for downstream analyses. N = 24,298,869 CpGs analyzed. The genome was split into 1Kb bins using the function makewindows from the bedtools^82^, package. DNA methylation signal was quantified by calculating the mean methylation on these 1Kb bin regions. Only bins with at least 3 CpGs covered at least 5X in all tested samples were used for downstream analyses. N= 2,807,124 bins analyzed.

Dynamic DNAm bins containing Gene Regulatory Elements (GREs) were identified by overlapping those bins with peaks of ATAC-seq, either located at a TSS (promoter) or at an H3K4me1 peak as previously described^26^.

### Analysis of DNA methylation arrays

IDAT raw data from the MethylationEPIC BeadChip 850k v2.0 microarrays was loaded using R to perform all the analyses, QC, and preprocessing steps using the package minfi (v1.42.0)^89^. Probes with low detection p-value (< 0.1), probes with a known SNP at the CpG site, and known cross-reactive probes were removed. For the resulting CpGs, beta values were calculated using minfi functions. DMPs were calculated using the limma package (v3.52.2)^84^ in R, adjusting by the Benjamini-Hochberg method. Only DMPs with adjusted p-values (FDR) < 0.05 and a differential of DNA methylation equal to or greater than 30% (Δβ ≥ 0.3) were selected for further analyses.

### RNA-seq analysis

Reads were mapped using STAR (v2.7.6)^90^. Gene expression was quantified using the function featureCounts from the package Subread (v2.0.3)^91^. Differentially expressed genes were detected using the R package DESeq2 (v1.36.0)^86^, applying P value < 0.05 filtering and log2FC filtering of +/-0.5. Transcription factors’ activities were inferred from expression values using Dorothea^40^.

### ChIP-seq analysis

For the ChIP-seq analyses, reads were trimmed using the TrimGalore package (v0.6.6) to remove adaptors and mapped using Bowtie2 (v2.4.4.1)^92^. Duplicated reads were removed with the tool MarkDuplicates from the package Picard (v3.1.0)^93^. Peaks were called using MACS2 (v2.2.5) ^94^, parameter (-q 0.05). Differential enrichment analysis was performed using the edgeR^95^, function from the diffBind package (v3.6.5) and filtering by FDR (0.05) and log2FC (2).

### Data availability

Data generated in this study are available in the Gene Expression Omnibus (GEO) database under accession numbers: WGBS-seq (GSE270886), dCas9-DNMT3A ChIP-seq (GSE270885), RNA-seq (GSE270876 and GSE270873) and MethylationEPIC BeadChip 850k v2.0 microarrays (GSE268284). ATAC-seq and CEBPA ChIP–seq datasets during human B-to-macrophage transdifferentiation used in the current study are available in the GEO database under accession number GSE131620. The H3K27ac, H3K4me1 and H3K4me3 ChIP–seq datasets used in this study are available in the ArrayExpress database under accession number E-MTAB-9010. The H3K27ac ChIP-seq datasets in mouse HSPCs and GMPs used in the current study are available in the GEO database under accession number GSE59636. The CEBPA ChIP-seq datasets in mouse HSPCs and GMPs used in the current study are available in the GEO database under accession number GSE43007.

## SUPPLEMENTARY INFORMATION

Supplementary Table 1. sgRNA sequences. Supplementary Table 2. RT-qPCR primer sequences.

Supplementary Table 3. Pyrosequencing primer sequences and genomic positions. Supplementary Table 4. ChIP-qPCR primer sequences.

## ACKNOWLEDGMENTS

We thank Dr. Thomas Graf and Dr. Bruno Di Stefano for critically reading the manuscript and all members of the Sardina group for stimulating scientific discussions. We thank Dr. X. Shawn Liu for sharing protocols for cellular transduction with the DNA methylation editors, Dr. Gregoire Stik for sharing the list of GREs in the human B to Macrophage transdifferentiation and the IJC Genomics Unit for their technical assistance with the processing of the MethylationEPIC arrays. This work was supported by the Grants PID2019-111243RA-I00 and PID2022-140376OB-I00 funded by MICIU/AEI/10.13039/501100011033 and FEDER/EU; the Worldwide Cancer Research (Grant reference 20-0269) and the Departament de Recerca i Universitats de la Generalitat de Catalunya (2021 SGR 01213). J.L.S. was funded by Instituto de Salud Carlos III through the project CP19/00176 (co-funded by the European Social Fund, “Investing in your future”). We thank CERCA Programme/Generalitat de Catalunya for institutional support.

## AUTHOR CONTRIBUTIONS

G.V. designed and performed the cell culture and molecular biology experiments, analyzed data, prepared figures, and edited the manuscript. A.V.L-R and A.L. designed and performed bioinformatic analyses and prepared figures. C.B. assisted G.V. with cell culture and molecular biology experiments. J.C. assisted G.V. with flow cytometry and immunofluorescence experiments. J.R-U and E.B. provided valuable advice and edited the manuscript. J.L.S. conceived the study, supervised the research, prepared figures, and wrote the manuscript with input from all co-authors.

---

**Extended Data Figure 1.**
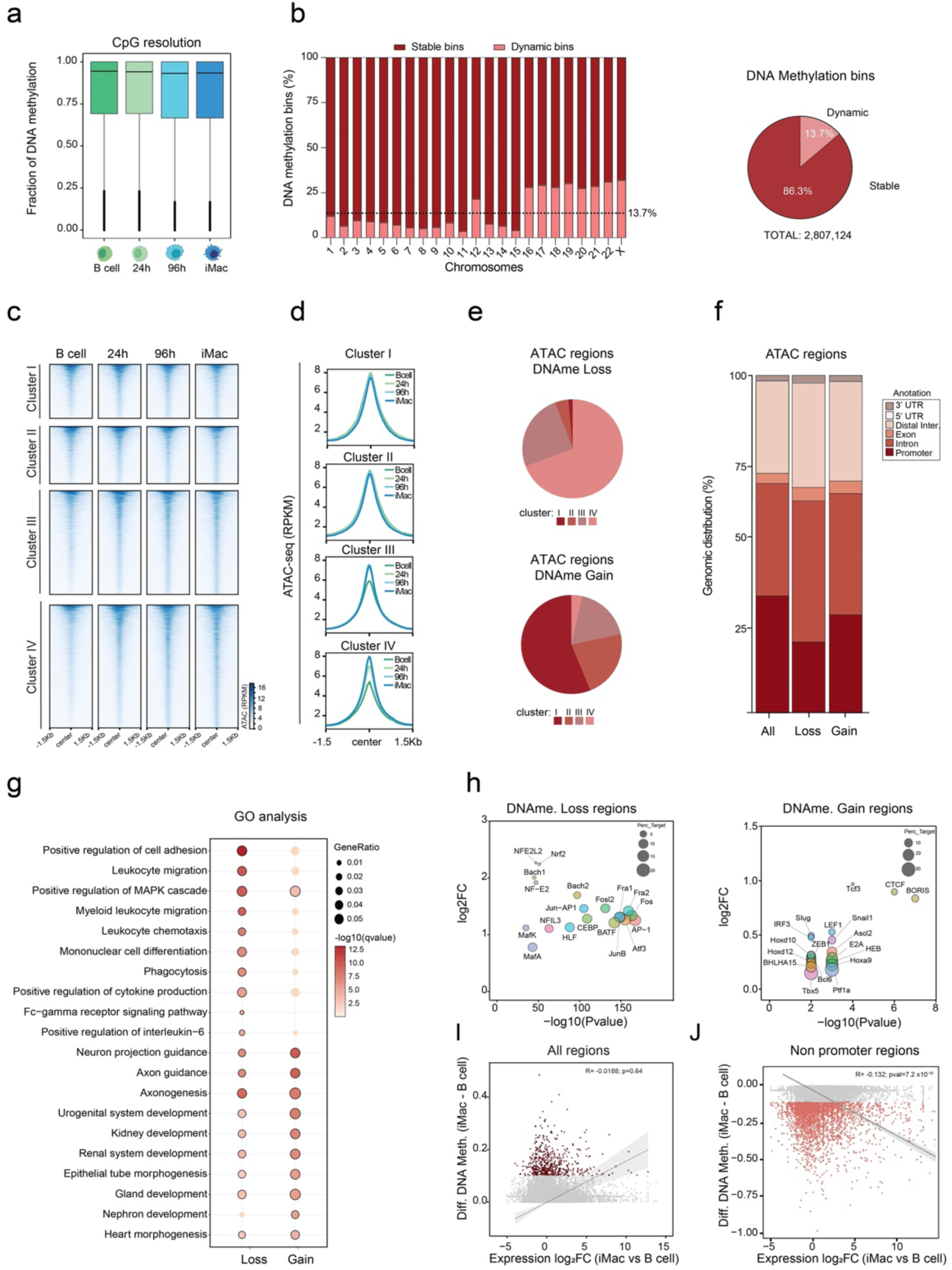
**(a)** Genome-wide DNAm levels at CpG resolution during transdifferentiation. Plots display the median (bar), interquartile range (box), and the 95% confidence interval (vertical bar). N (number of CpGs analyzed) = 24,298,869 **(b) Left:** Percentage of DNAm stable and dynamic 1kb-bins observed during transdifferentiation across chromosomes. The dashed line represents the genome-wide percentage of dynamic bins (13.7%); **Right:** Pie chart showing the genome-wide percentage of DNAm stable and dynamic 1kb-bins observed during transdifferentiation. N (number of bins analyzed) = 2,807,124. **(c)** Genomic heatmaps showing ATAC-seq signal along transdifferentiation at the DNAm clusters in Fig. 1c. ATAC-seq data from GSE131620^26,28^. **(d)** Average plots of ATAC-seq signal along transdifferentiation at the DNAm clusters in Fig. 1c. **(e)** Piecharts depicting the proportion of ATAC+ DNAm loss and gain regions in Fig. 1e belonging to the methylation clusters in Fig. 1c. **(f)** Distribution of ATAC+ DNAm loss and gain regions in iMacs along the different genomic features analyzed (promoters, introns, exons, distal intergenic, 5’ UTR and 3’UTR regions). All ATAC+ regions analyzed (All) are shown as a distribution control. **(g)** Balloon plot depicting the Gene Ontology (GO) enrichment analysis for the genes associated with the ATAC+ DNAm loss and gain regions in iMacs (related to Fig. 1d). 10 highly significant over-represented biological processes (BP) terms for each category are plotted. Ratio of genes of interest over all unique genes (GeneRatio) and - log10(qvalue) are shown. Significant terms (qvalue < 0.05) are highlighted with a black stroke. **(h)** Bubble plot showing the output of transcription factor enrichment analyses (by HOMER2) at ATAC+ DNAmth loss **(left)** or gain **(right)** regions in iMacs. X-axis reports the –log10(P-value); Y-axis depicts the log2FC; the circle size is related to the percentage of target sequences containing the motif. **(i)** Plot showing the correlation between the step changes (iMac-Bcells) in DNAm and gene expression at iMac’s chromatin accessible regions (ATAC+ peaks). Grey dots: ATAC+ regions not gaining at least 10% in DNAm. Dark red dots: ATAC+ regions gaining at least 10% in DNAm in iMacs. RNA-seq data from GSE140528^26,28^. **(j)** Plot showing the correlation between the step changes (iMac-Bcells) in DNAm and gene expression at iMac’s chromatin accessible regions (ATAC+ peaks) non-overlapping annotated promoters. Grey dots: Regions not losing at least 10% in DNAm. Light red dots: Regions losing at least 10% of DNAm.

**Extended Data Figure 2.**
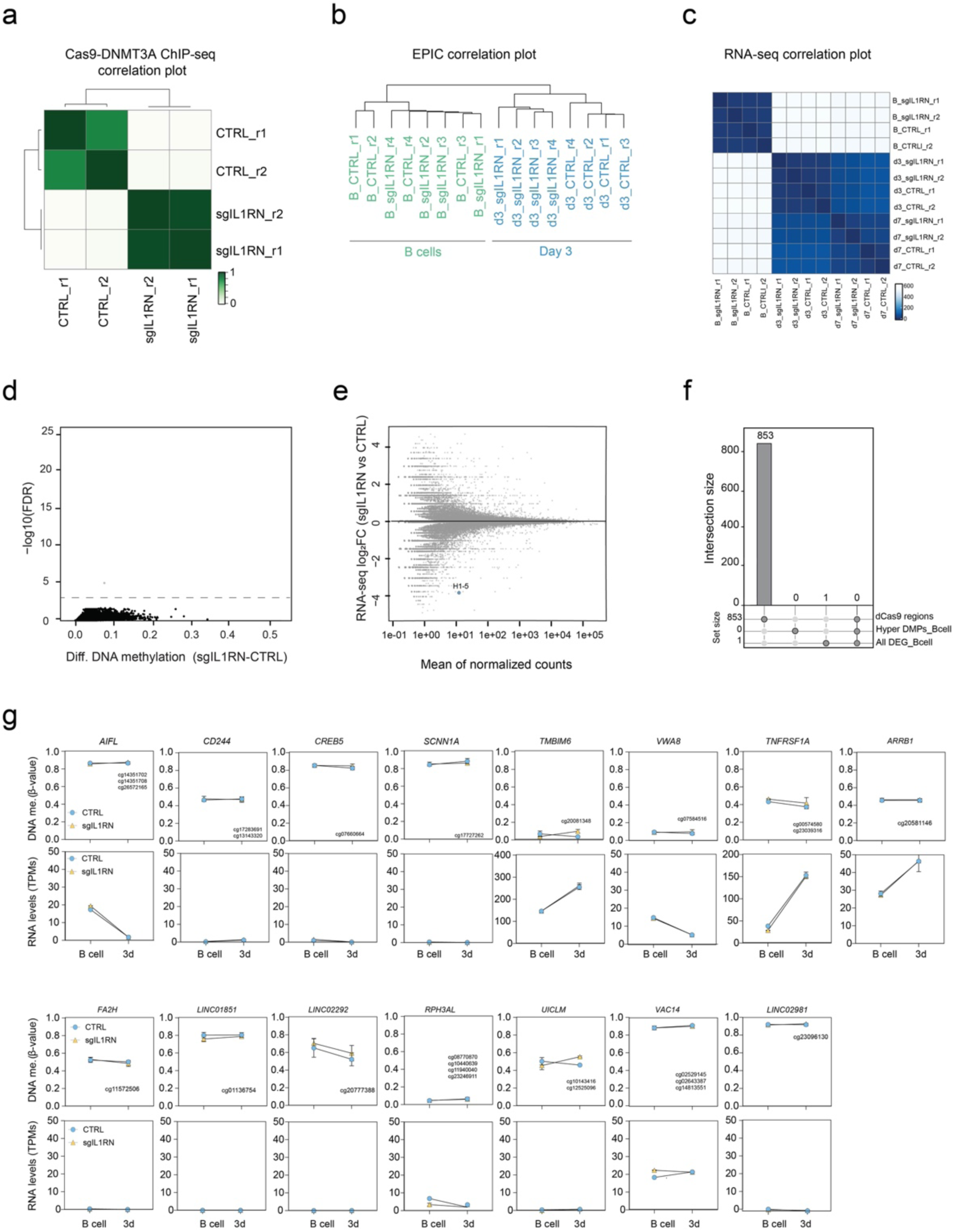
**(a)** Correlation heatmap showing the correlation (r) values for the Cas9 ChIP-seq experiments in dCas9-DNMT3A CTRL and sgIL1RN B cells. **(b)** Unsupervised clustering of the transdifferentiation samples subjected to the MethylationEPIC BeadChip 850k v2.0 microarrays. **(c)** Correlation heatmap showing the correlation (r) values between the RNA-seq samples during transdifferentiation. Scale bar represents the range of the correlation coefficients (r) displayed. **(d)** Scatter plot showing differentially hypermethylated CpGs in dCas9-DNMT3A sgIL1RN B cells compared to CTRL B cells. Black dots indicate non-significantly hypermethylated CpG positions (FDR > 0.05). The grey dot indicates a significantly hypermethylated position (FDR < 0.05). No significantly hypermethylated CpG positions gaining ≥ 30% of DNAm (FDR < 0.05, Δβ < 0.3) were observed. The dashed line indicates the FDR < 0.05. **(e)** MA plot showing differentially expressed genes (DEGs) in dCas9-DNMT3A sgIL1RN B cells compared to CTRL B cells. The blue dot indicates a significant DEGs (FC>1, p<0.05, n=3). **(f)** Upset plot depicting the intersection of the significantly dCas9-DNMT3A bound regions (in Fig. 2f), the significantly hypermethylated ⃤β ≥ 0.3 CpG positions (in d), and the significant associated DEGs (in e) in dCas9-DNMT3A sgIL1RN compared to CTRL B cells. No candidate intersects the 3 datasets. **(g)** Plots showing the DNAm (upper panels) and transcriptional (bottom panels) dynamics of top dCas9-DNMT3A bound regions (FC>3.5, p<0.05) identified in Fig. 2g. Unpaired two-tailed Student’s t-test, n=4 (DNAm) n=2 (RNA-seq) per group, mean ± s.e.m, not significant (p< 0.05).

**Extended Data Figure 3.**
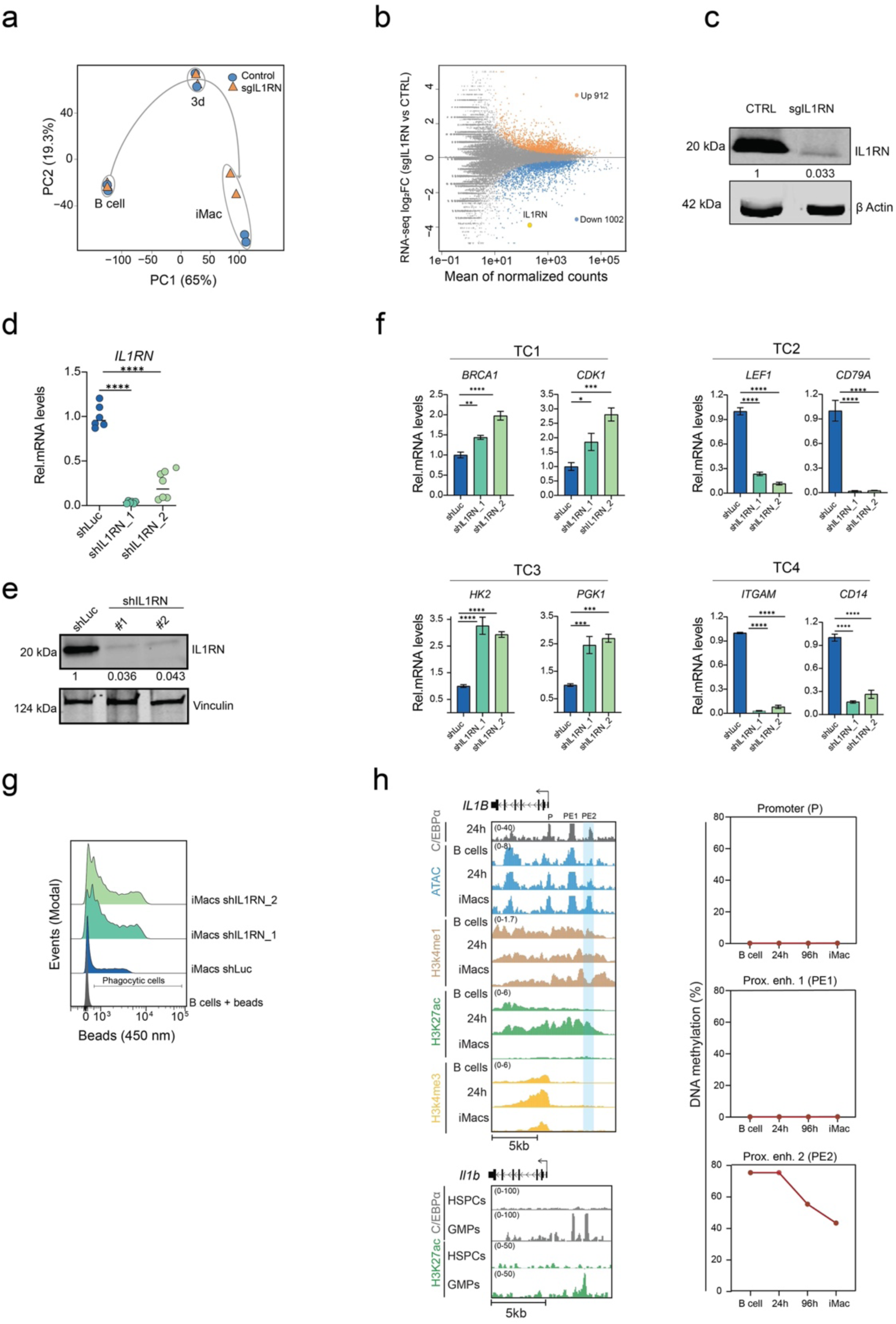
**(a)** Principal Component Analysis (PCA) displaying the transcriptomic dynamics of dCas9-DNMT3A sgIL1RN and CTRL cells during transdifferentiation. **(b)** MA plot showing differentially expressed genes (DEGs) in dCas9-DNMT3A sgIL1RN iMacs compared to dCas9-DNMT3A CTRL iMacs. Orange dots indicate upregulated genes (FC 1 and p<0.05, n= 912). Blue dots and the yellow dot (*IL1RN*) indicate downregulated genes (FC -1 and p<0.05, n= 1002). **(c)** Representative western blot image of IL1RN protein in dCas9-DNMT3A sgIL1RN and CTRL iMacs. IL1RN levels were normalized to β-Actin levels and expressed as a fold change over CTRL iMacs. **(d)** qRT-PCR analysis of IL1RN expression in iMacs harboring shRNAs targeting the luciferase gene (shLuc, control) or the *IL1RN* gene (shIL1RN). Values were normalized against *B2M* expression. Two-way ANOVA with Dunnett’s post-hoc test, n=6, mean ± s.e.m., (****p < 0.0001). **(e)** Representative western blot image of IL1RN protein in shIL1RN and shLuc iMacs. IL1RN levels were normalized to vinculin protein levels and expressed as a fold change over shLuc iMacs. **(f)** qRT-PCR analysis in shIL1RN and shLuc iMacs of selected genes from the transdifferentation clusters shown in Fig. 3b. Values were normalized against *B2M* expression. Two-way ANOVA with Dunnett’s post-hoc test, n=3, mean ± s.e.m., (*p < 0.05; **p < 0.01; ***p < 0.001; ****p < 0.0001). **(g)** Flow cytometric analysis showing the Mean Fluorescence Intensity (MFI) for the myeloid markers (CD11b and CD14) in shIL1RN and shLuc iMacs. Two-way ANOVA with Dunnett’s post-hoc test, n=3, mean ± s.e.m., (****p < 0.0001). **(h) Top**: Genome browser snapshot at the human *IL1B* locus showing signal for C/EBPα ChIP-seq at 24h and chromatin accessibility (by ATAC-seq), H3K4me1, H3K27ac and H3K4me3 ChIP-seq signals in B cells, 24 hours and in iMacs. At 24h, C/EBPα binds 3 GREs within the *IL1B* locus: the promoter (P) and 2 proximal enhancers (PE1-2). The blue-shaded highlight indicates a DNA demethylation event at PE2 (as identified in Fig. 1e). DNAm kinetics at the 3 GREs are illustrated on the right panels. **Bottom:** Genome browser snapshot at the mouse *Il1b* locus showing signal for C/EBPα and H3K27ac ChIP-seq in murine Hematopoietic Stem Progenitor cells (HSPCs) and Granulo-Monocyte Progenitors (GMPs). Data were taken from: H3K27ac, H3K4me1 and H3K4me3 ChIP-seq during human transdifferentiation (ArrayExpress: E-MTAB-9010); C/EBPα ChIP-seq in murine HSPCs and GMPs (GEO: GSE43007)^71^ ChIP-seq signal in murine HSPCs and GMPs (GEO: GSE59636)^72^.

**Extended Data Figure 4.**
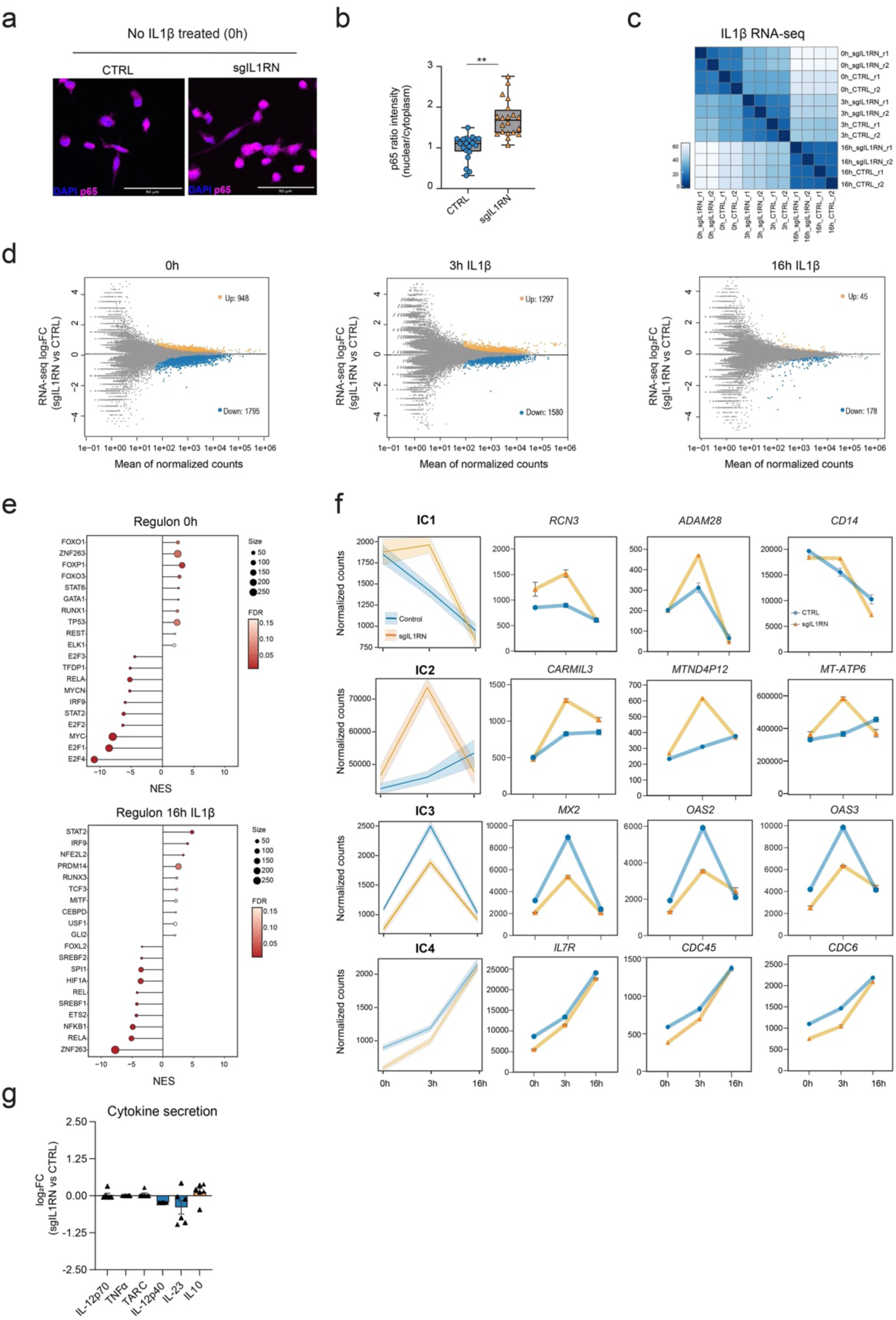
**(a)** Representative IF showing p65 (red) and DAPI (blue) signals in dCas9-DNMT3A sgIL1RN and CTRL untreated (0h) iMacs. **(b)** Quantification of p65 nuclear versus cytoplasmic localization signal in dCas9-DNMT3A sgIL1RN and CTRL iMacs. Unpaired two-tailed Student’s t-test, n=20 cells per group, mean s.e.m., (**p< 0.01). **(c)** Correlation heatmap showing the correlation (r) values between the RNA-seq samples during IL1β treatment. Scale bar represents the range of the correlation coefficients (r) displayed. **(d)** MA plots showing differentially expressed genes (DEGs) during IL1β treatment in dCas9-DNMT3A sgIL1RN iMacs compared to CTRL iMacs. Orange dots indicate upregulated genes (FC 1 and p<0.05). Blue dots (*IL1RN*) indicate downregulated genes (FC -1 and p<0.05). DEGs: (0h) up=948, down=1795; (3h IL1β) up=1297, down=1580; (16h IL1β) up=45, down=178. **(e)** Lollipop plot depicting the TF activity predicted from mRNA expression of target genes with DoRothEA v2.0^40^ in untreated (0h) dCas9-DNMT3A sgIL1RN and CTRL iMacs (upper panel) and treated with IL1β for 16h (bottom panel). Top 10 most significant (FDR < 0.05) transcriptional target genes from positive and negative normalized enrichment are shown. Lollipop size indicates the total number of differentially expressed genes regulated by each transcriptional regulon. **(f) Left:** Quantification of RNA signal at the IL1β-treatment clusters (IC1-4) in Fig. 4e.; **Right:** expression dynamics of representative genes from the clusters in dCas9-DNMT3A sgIL1RN and CTRL iMacs. **(g)** Normalized levels of secreted cytokines in dCas9-DNMT3A sgIL1RN iMacs treated for 24h with IL1β. One-way ANOVA with Dunnett’s post-hoc correction, n=6, mean, not significant (p>0.05).

