## Supplementary information for "Modulating immune cell fate and inflammation through CRISPR-mediated DNA methylation editing"

**Supplementary Table 1:** Oligonucleotides used to construct sgRNAs targeting with dCas9 methylation editing tool**.**

| | **Forward (5' to 3')** | **Reverse (5' to 3')** |
|---|---|---|
| ILsgRNA_H1 | TCCCACTCCATTGCGACACTTAGTG | CACTAAGTGTCGCAATGGAG |
| ILsgRNA_mU6 | TTGTTTGCTTCTCGCAGTGGGGCAGGG | CCCTGCCCCACTGCGAGAAG |
| ILsgRNA_ hU6 | CACCGAGCTTGGGTGAGTGACTATT | AATAGTCACTCACCCAAGCT |
| ILsgRNA_7SK | CCTCGCCACAACTCTGGGCCCGCAA | TTGCGGGCCCAGAGTTGTGG |
| CTRLsgRNA_H1 | TCCCACCTAAGGTTAAGTCGCCCTC | AAACGATTCCAATTCAGCGGGAGT |
| CTRLsgRNA_mU6 | TTGTTTGGCCCCCGGGGGAAAAATTT | AAACCCGGGGGCCCCCTTTTTAAACAA |
| CTRLsgRNA_ hU6 | CACCGTAGTACTTTCAAGAGTCCA | AAACTATCATGAAAGTTCTCAGGTC |
| CTRLsgRNA_7SK | CCTCGCACTACCAGAGCTAACTCA | AAACCGTGATGGTCTCGATTGAGT |

**Supplementary Table 2**: Primers used for RT-qPCR to determine the expression levels in sgIL1RN and sgCTRL cells.

| | **Forward (5' to 3')** | **Reverse (5' to 3')** |
|---|---|---|
| IL1RN | IDT Cat #228354241 | CTGCATTGTTTTGCCAGTGT |
| HPRT | GACCAGTCAACAGGGGACAT | CTGCATTGTTTTGCCAGTGT |
| B2M | AGGCTATCCAGCGTACTCCA | TCAATGTCGGATGGATGAAA |
| BRCA1 | CTGCTCTGGGTAAAGTTCATTGG | TAAAGGACACTGTGAAGGCCC |
| CDK1 | CACTTGGCTTCAAAGCTGGC | TGGGTATGGTAGATCCCGGC |
| LEF1 | ATTCTTGGCAGAAGGTGGCA | GCAGCTGTCATTCTTGGACC |
| CD79A | CCTTAGTCATATTCCCCCAG | TTTAGAGGGAAGAAGAGTGG |
| HK2 | GCTCAACCATGACCAAGTGC | AACTCTCCGTGTTCTGTCCC |
| PGK1 | CTGGGCAAGGATGTTCTGTT | CACATGAAAGCGGAGGTTCT |
| ITGAM | GGGGTCTCCACTAAATATCTC | CTGACCTGATATTGATGCTG |
| CD14 | GATTACATAAACTGTCAGAGGC | TCCATGGTCGATAAGTCTTC |

**Supplementary Table 3.** List of genomic regions of the CpGs located at *IL1RN* promoter analyzed by pyrosequencing. CpG code corresponds at **Fig2a** and **2c**.

| CpG code | Genomic position hg38 |
|---|---|
| #1 | chr2:113,127,510-113,127,511 |
| #2 | chr2:113,127,517-113,127,518 |
| #3 | chr2:113,127,539-113,127,540 |
| #4 | chr2:113,127,589-113,127,590 |

Primers used in this study for the PCR amplification of *IL1RN* promoter after bisulfite conversion for pyrosequencing analysis.

| | Forward (5' to 3') | Reverse-bio (5' to 3') | Region (5' to 3') hg38 |
|---|---|---|---|
| IL1RN promoter | AGTGGGGTTGAAAGTGACAAC | CAGAATGGAAATCTGCAGAGGCCTC | chr2:113,127,448-113,127,645 |

Primers used in this study for sequencing by pyrosequencing and assess the methylation levels of target CpG sites.

| | Forward (5' to 3') |
|---|---|
| S1 IL1RN | GAAATGCGAGGAGGGTATTTCCGCTTCTCG |
| S2 IL1RN | CGCTTCTCGCAGTGGGGCAGGGTGGCAGACGC |

**Supplementary Table 4**. Primers used for ChIP-qPCR to determine the enrichment of Cas9 in negative control region and the *IL1RN* promoter.

| | Forward (5' to 3') | Reverse (5' to 3') |
|---|---|---|
| IL1RN –5kb TSS | GCAGTCGGGGTTGGGGTAA | ACTCAGGCTAGCAGAAACCAA |
| IL1RN promoter | GGAGGGTATTTCCGCTTCTC | GCCTCTGCAGATTTCCATTC |
